# A pan-cancer analysis of microRNA tissue specificity and its association with dysregulation

**DOI:** 10.64898/2026.06.01.729194

**Authors:** Aly Ismailov, Alexey Belogurov, Alena Evpak, Maria Poptsova

## Abstract

MicroRNAs are frequently dysregulated in cancer, yet how their tissue-specificity is remodeled during malignant transformation remains poorly characterized. Here we systematically quantified the tissue-specificity of miRNAs across normal (GTEx) and tumor (TCGA) tissues using the Tau index, and compared its distribution between healthy and cancerous states. To robustly define dysregulation, we combined two independent analyses: a binomial test over per-project differential expression across 17 matched normal tissues within TCGA cohort, and a TCGA–GTEx pan-tissue comparison of mean expression. The change in specificity (ΔTau) separated up- from down-regulated miRNAs, showing moderate agreement with the binomial signal and a strong correlation with the expression-based contrast. Finally, we identified 6 miRNAs that lose tissue-specificity upon transformation while remaining consistently upregulated (miR-519a-5p, miR-512-3p, miR-522-3p, miR-105-5p, miR-935, miR-1269a). Functional analysis of experimentally validated targets showed significant enrichment for converging on core oncogenic programs for miR-512-3p, miR-105-5p and miR-935, such as apoptosis and cellular-stress regulation, TP53, FoxO, PI3K–Akt/mTOR signaling, immune modulation. Collectively, integrating specificity dynamics with dysregulation evidence pinpoints candidate miRNAs with coordinated, cancer-relevant regulatory roles and highlights those with favorable tissue specificity profiles for therapeutic targeting.

## 1. Introduction

MicroRNAs (miRNAs) are small non-coding RNAs that act as key post-transcriptional regulators of gene expression and play essential roles in the maintenance of cellular identity, differentiation, and tissue homeostasis. By fine-tuning the expression of multiple target genes simultaneously, miRNAs coordinate diverse biological processes, including development, proliferation, apoptosis, and immune responses. Dysregulation of miRNA expression is a well-established hallmark of cancer and contributes to numerous oncogenic processes, including uncontrolled proliferation, invasion, metastasis, angiogenesis, immune evasion, and cellular plasticity [1].

Over the past two decades, extensive studies have identified tumor-associated miRNAs and established their potential utility as diagnostic biomarkers, prognostic indicators, and therapeutic targets [1]. However, most pan-cancer investigations have focused primarily on differential expression analysis, emphasizing whether specific miRNAs are upregulated or downregulated in tumors relative to normal tissues [2-4]. While this approach has yielded important insights, it does not capture important aspect of miRNA biology, that is tissue specificity. In normal tissues, tissue-specific miRNAs help stabilize lineage-associated transcriptional states and suppress alternative differentiation programs, thereby maintaining cellular phenotype and functional specialization [5-7]. Disruption of these regulatory constraints may represent a critical feature of malignant transformation, enable dedifferentiation and increase cellular plasticity. Nevertheless, the extent to which loss of tissue specificity contributes to recurrent pan-cancer miRNA deregulation remains poorly understood.

In this study, we performed an integrative analysis of two major cohorts: healthy tissues from the Genotype-Tissue Expression (GTEx) project and tumor samples from TCGA, TARGET, CGCI, and CPTAC, hereafter collectively referred to as the TCGA cohort for simplicity. Tissue specificity was quantified using the Tau score across tissues in both cohorts. To identify miRNAs associated with malignant transformation, we applied two complementary analytical approaches. First, differential expression analysis was performed within individual cancer projects by comparing tumor samples with matched normal tissues where available. Second, miRNA expression patterns and tissue specificity profiles were compared between the GTEx and TCGA cohorts. The results obtained from both approaches were integrated to identify robust miRNA signatures consistently associated with malignant transformation. This analysis revealed 6 miRNAs that lost their tissue specificity upon malignant transformation, shifting from tissue-restricted expression patterns in healthy tissues to widespread activation across tissues in cancer samples. Enrichment analysis of experimentally validated targets of these miRNAs supported their involvement in cancer-related biological processes, highlighting their potential as candidates for antagomir-based therapeutic strategies due to their restricted expression in normal tissues and broad activation across tumor types.

### 2. Materials and methods

#### 2.1. Data preprocessing and inclusion criteria

MiRNA expression profiles from the healthy (GTEx) [8] and tumor (TCGA, TARGET, CGCI, CPTAC) tissues [9] databases were used in this study. During preprocessing, samples were grouped according to normal tissue types and cancer tissue types. Only tissues represented by at least 10 biological samples were included in the analysis. To ensure comparability between normal and tumor datasets, tissue categories were harmonized based on the closest possible correspondence in both tissue type and number of represented tissues. As a result, 26 tissue types from GTEx and 25 tissue types from TCGA were included in the comparative analysis.

MiRNAs with a maximum tissue expression level below 20 counts per million (CPM) were filtered out separately for the GTEx and TCGA datasets. Subsequent analyses were restricted to miRNAs detected in both datasets. Following these filtering steps, the initial sets of 2,011 miRNAs from TCGA and 827 miRNAs from GTEx were reduced to 394 miRNAs retained for further analysis.

#### 2.2. Tissue specificity analysis (Tau index)

The degree of tissue-specific expression for each miRNA was quantified using the Tau (τ) tissue specificity index [10, 11]. Tau values were calculated based on the log2-transformed mean miRNA expression levels across tissue groups after CPM normalization, with a pseudocount of 1 added to avoid zero values. The Tau index was calculated according to the following formula:

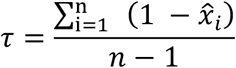

where *n* denotes the total number of groups, and 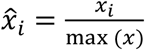 represents the normalized expression of a given miRNA in the *i*-th group relative to its maximum expression value across all groups. Tau values range from 0 (ubiquitous expression) to 1 (strict tissue specificity). The index was calculated independently for the healthy and cancer cohorts. Specificity-based classification. Using a threshold of τ > 0.8, all analyzed miRNAs were classified into four functional categories according to their tissue specificity patterns in normal and tumor tissues:

1. Normal-tissue-specific miRs. miRNAs exhibiting tissue specificity only in normal tissues (τ_*GTEx*_ ≥0.8, τ_*TCGA*_ <0.8).
2. Cancer-tissue-specific miRs. miRNAs exhibiting tissue specificity exclusively in cancer tissues (τ_*TCGA*_ ≥0.8,τ_*GTEx*_ <0.8).
3. Tissue specific miRs. miRNAs retaining high tissue specificity in both datasets (τ ≥ 0.8in both cohorts).
4. Non-specific miRs. miRNAs exhibiting non-specific expression patterns in both normal and tumor conditions (τ <0.8).

#### 2.3. Differential expression analysis within TCGA cohort and binomial test

To identify dysregulated miRNAs, expression levels between tumor and normal tissues were compared within each cancer project containing paired controls in TCGA dataset. A total 17 projects were included, meeting the criterion of ≥10 normal samples and ≥10 tumor samples. Differential expression analysis was performed using the PyDESeq2 library [12]. Model parameters were estimated using DeseqDataSet and DeseqStats with refit_cooks=True to minimize the influence of outliers. The comparison was performed for tumor vs normal, and results were considered statistically significant at an adjusted p-value < 0.01 and |log_2_FC| > 1.

To assess whether individual miRNAs exhibited a significant bias toward upregulation or downregulation across cancer projects, we applied a one-sided exact binomial test. For each miRNA, the number of significantly upregulated (*n_up_*) and downregulated (*n*_*down*_) instances was counted based on the thresholds defined above (with the direction determined by the sign of the log_2_FC), and the total number of dysregulated cases was defined as: *n*_*total*_ = *n_up_* + *n*_*down*_. Non-significant cases were intentionally excluded from *n*_*total*_ to maintain statistical power and account for heterogeneous nature of cancers. Under the null hypothesis, upregulation and downregulation were assumed to be equally likely among significantly dysregulated cases (H_0_ : 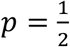 ; where p denotes the probability of upregulation). Two one-sided exact binomial tests were performed: *alternative = “greater”* to test whether upregulated events occurred more frequently than expected by chance, and *alternative = “less”* to test whether downregulated events occurred more frequently than expected by chance. Statistical significance was defined as p < 0.05.

#### 2.4. Comparison of mean miRNA expression between GTEx and TCGA tissues

As an additional criterion, mean miRNA expression levels were compared between tissues from the GTEx and TCGA cohorts. Differences in miRNA expression across tissues between the two cohorts were assessed using the Mann–Whitney rank-sum test applied to log2(CPM + 1) transformed expression values, followed by Benjamini–Hochberg correction for multiple testing (false discovery rate, FDR). The effect size was estimated using log2 fold change calculated from the mean expression values between the two cohorts. Statistical significance was defined as FDR < 0.01 combined with an absolute log_2_FC value greater than 1 (|log2FC| > 1).

## 3. Results

### 3.1. Tissue specificity of miRNAs in cancer and healthy cohorts

To enable a comparable analysis between the GTEx and cancer cohorts, miRNA expression profiles were stratified according to tissue of origin, resulting in 25 tissue groups in the cancer cohort and 26 corresponding tissue groups in the GTEx dataset (Fig. 1A, B). The overall distribution of tissue specificity scores (Tau) demonstrated comparable patterns between normal and tumor tissues, with median Tau values of 0.44 and 0.38 in GTEx and TCGA cohorts, respectively (Fig. 1C). Using a Tau threshold of >0.8, miRNAs were classified into four categories according to their tissue specificity patterns: tissue-specific miRNAs, cancer-tissue-specific miRNAs, normal-tissue-specific miRNAs, and non-specific miRNAs (Fig. 1D). Analysis of changes in tissue specificity (ΔTau) revealed a modest trend toward loss of tissue specificity in cancer tissues compared with healthy tissues (Fig. 1E).

**Fig 1.**
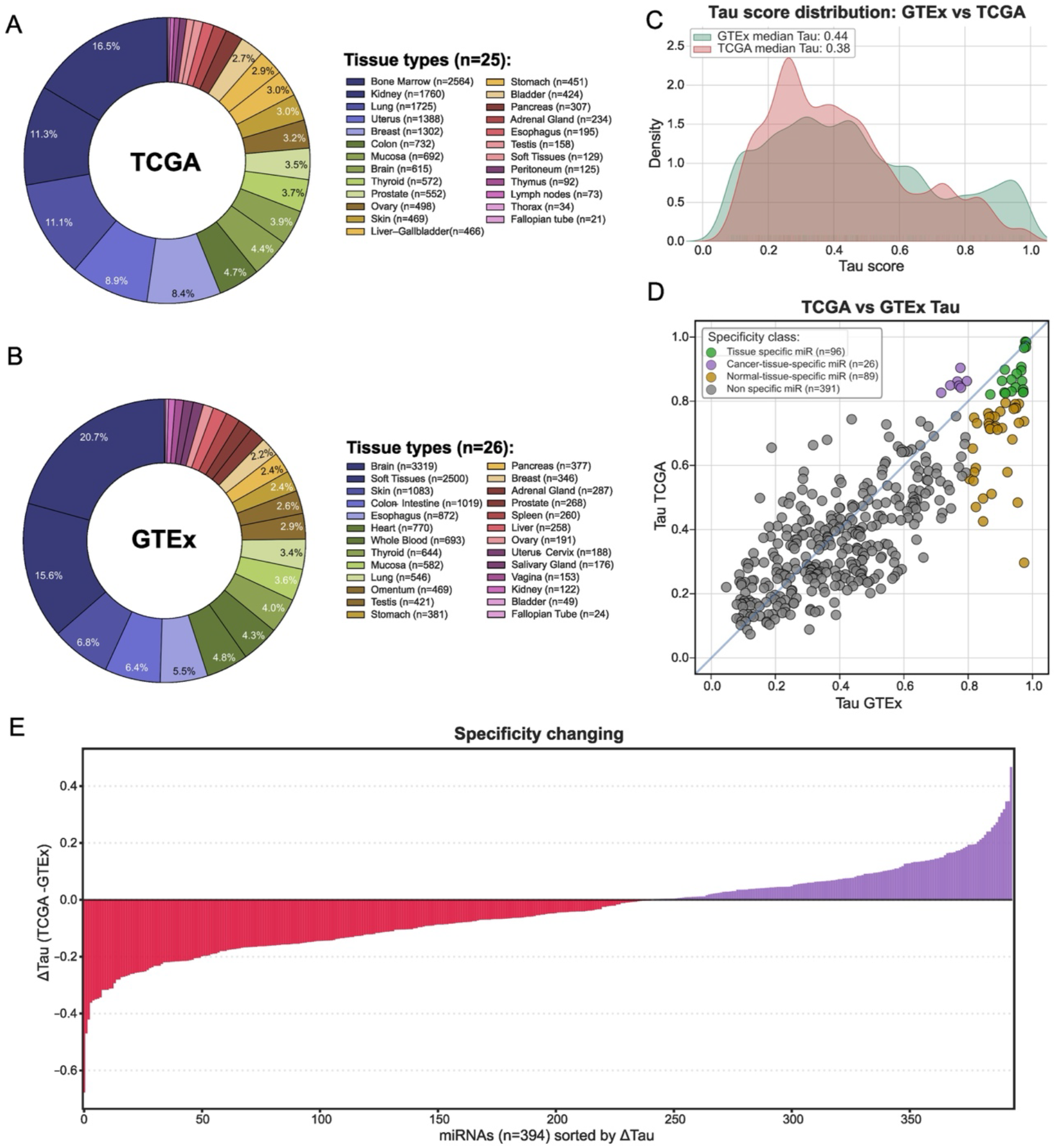
Overview of the datasets and tissue specificity analysis. (A) Distribution of healthy tissue types in the GTEx cohort. (B) Distribution of cancer tissue types in the TCGA cohort. (C) Distribution of tissue specificity (Tau) scores across tissues in the GTEx and TCGA cohorts. (D) Scatter plot of tissue specificity (Tau) scores for individual miRNAs in the GTEx (x-axis) and TCGA (y-axis) cohorts. Using the commonly applied Tau threshold of 0.8, miRNAs were classified into four categories: tissue-specific (green), cancer-tissue-specific (purple), normal-tissue-specific (yellow), and non-specific (gray). (E) Distribution of ΔTau values across miRNAs, illustrating changes in tissue specificity between normal and cancer cohorts. Red lines indicate miRNAs with negative ΔTau values (decreased tissue specificity in cancer), whereas purple lines indicate miRNAs with positive ΔTau values (increased of tissue specificity in cancer) among the 394 miRNAs common to both cohorts.

### 3.2. Dysregulated miRNAs in cancer

Differential expression analysis was performed independently across 17 cancer projects with available matched tumor and normal samples (Fig 2A). Volcano plots for each cohort are provided in the Supplementary Fig S1, and the complete differential expression results are available in the Supplementary Table S1. According to the binomial test (See Methods), a large fraction of miRNAs (n = 311, 78.9%) did not show a significant enrichment toward either up- or downregulation among significant differential expression events. Among all analyzed miRNA, 14.6% (n = 57) were classified as recurrently upregulated, while 6.6% (n = 26) were classified as recurrently downregulated across cancer types (Fig 2B).

**Fig 2.**
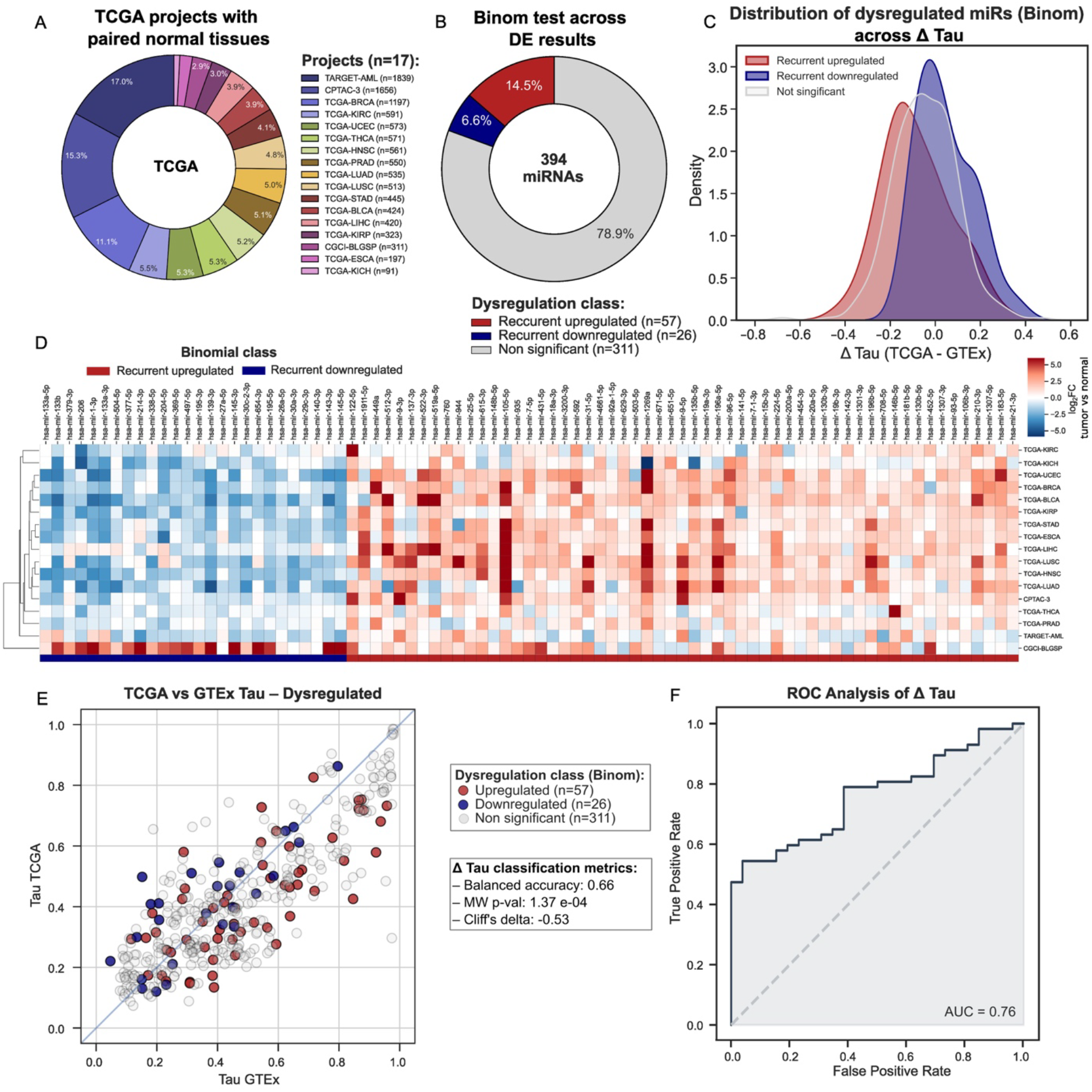
Identification of recurrently dysregulated miRNAs and their association with changes in tissue specificity. (A) Distribution of cancer projects within the TCGA cohort with available matched normal tissues. (B) Results of the directed binomial test following differential expression (DE) analysis across 17 cancer cohorts. Among 394 analyzed miRNAs, 311 (78.9%) did not demonstrate consistent dysregulation across cancer cohorts, whereas 57 miRNAs (14.5%) were recurrently upregulated and 26 miRNAs (6.6%) were recurrently downregulated. (C) Distribution of ΔTau values for dysregulated miRNAs. Recurrently upregulated miRNAs showed a shift toward negative ΔTau values, indicating reduced tissue specificity in cancer, whereas recurrently downregulated miRNAs tended toward positive ΔTau values. (D) Heatmap showing log2 fold-change values for 83 significantly dysregulated miRNAs across 17 cancer cohorts included in the differential expression analysis. (E) Scatter plot comparing tissue specificity scores (Tau) between GTEx and cancer cohorts. Each point represents an individual miRNA and is colored according to its recurrent differential expression status (upregulated, downregulated, or non-significant). The blue diagonal represents equal tissue specificity between normal and cancer cohorts, miRNAs below the diagonal exhibit reduced tissue specificity in cancer (negative ΔTau), whereas miRNAs above the diagonal exhibit increased tissue specificity (positive ΔTau). ΔTau showed moderate discrimination between recurrently upregulated and downregulated miRNAs (Balanced accuracy = 0.66, MW U test P = 1.37 × 10^-4^; Cliff’s δ = −0.53). (F) Receiver operating characteristic (ROC) curve evaluating the ability of Δtau to discriminate between recurrently upregulated and recurrently downregulated miRNAs. ΔTau demonstrated moderate discriminatory performance between the two classes (ROC-AUC = 0.76).

The distribution of dysregulated miRNAs along the ΔTau axis showed a modest shift of upregulated miRNAs toward negative ΔTau values, indicating a tendency to lose tissue specificity, whereas downregulated miRNAs were shifted toward positive ΔTau values, reflecting increased tissue specificity (Fig. 2C). The structure of dysregulation changes across TCGA projects is further illustrated in the heatmap (Fig 2D), which includes only statistically significant events (|log_2_FC| > 1, FDR < 0.01 for differential expression and p < 0.05 for the binomial test). Notably, the Burkitt Lymphoma Genome Sequencing Project (CGCI-BLGSP) shows a distinct inverted expression pattern. In this cohort, most miRNAs that are typically downregulated across other cancer types show significant upregulation (Fig 2D). ΔTau values differed significantly between recurrently upregulated and downregulated miRNAs (Mann–Whitney U test, P = 1.37 × 10^-4^), showing a moderate effect size (Cliff’s δ = −0.53). ΔTau demonstrated meaningful discriminatory ability between these two classes (ROC-AUC = 0.76). Using ΔTau = 0 as a threshold, the classification achieved a balanced accuracy of 0.66 (Fig. 2C).

As an additional analysis of miRNA dysregulation, we compared miRNA expression between the TCGA and GTEx cohorts using the Mann–Whitney U test (FDR < 0.01, |log_2_FC| > 1). This comparison identified a substantially larger number of differentially expressed miRNAs than the recurrent dysregulation analysis: 147 miRNAs were significantly upregulated in the TCGA cohort, 91 were significantly upregulated in the GTEx cohort, and 156 showed no significant differential expression (Fig. 3A). In contrast to the recurrent dysregulation analysis, this comparison revealed a much stronger separation of dysregulation classes along the ΔTau axis. miRNAs significantly upregulated in the TCGA cohort were predominantly shifted toward negative ΔTau values, indicating a loss of tissue specificity in cancer, whereas miRNAs upregulated in the GTEx cohort were enriched for positive ΔTau values, reflecting increased tissue specificity in normal tissues. In contrast, non-significant miRNAs were centered around ΔTau = 0 (Fig. 3B). This pattern was further supported by classification metrics. Using a biologically defined threshold of ΔTau = 0, ΔTau demonstrated excellent discrimination between miRNAs upregulated in the TCGA and GTEx cohorts, achieving a balanced accuracy of 0.9. The difference in ΔTau distributions was highly significant (Mann–Whitney U test, P = 1.9 × 10^-34^) with a very large effect size (Cliff’s δ = −0.95), while the overall discriminatory performance reached a ROC-AUC of 0.97 (Fig. 3C, D). This strong discriminatory performance might be explained by the relationship between expression changes and tissue specificity dynamics. A significant correlation between log_2_FC and ΔTau (Pearson ρ = −0.7), indicated that miRNAs with increased expression in cancer tissues tend to exhibit reduced tissue specificity (Fig. 3E, F). This relationship is not unexpected, as increased tissue specificity is generally accompanied by a decrease in the overall mean expression level across tissues, whereas reduced tissue specificity reflects broader expression patterns and consequently higher mean expression. Conversely, miRNAs with higher expression in normal tissues generally showed increased tissue specificity in the cancer cohort. Importantly, this strong coupling between expression changes and tissue specificity alterations suggests that mean expression differences between TCGA and GTEx, as assessed by the Mann–Whitney U test, may partly reflect shifts in tissue specificity rather than cancer-specific regulatory alterations alone. Therefore, miRNAs supported by both approaches, recurrent dysregulation across TCGA paired projects and consistent mean expression shifts between TCGA and GTEx were considered high-confidence dysregulated candidates.

**Fig 3.**
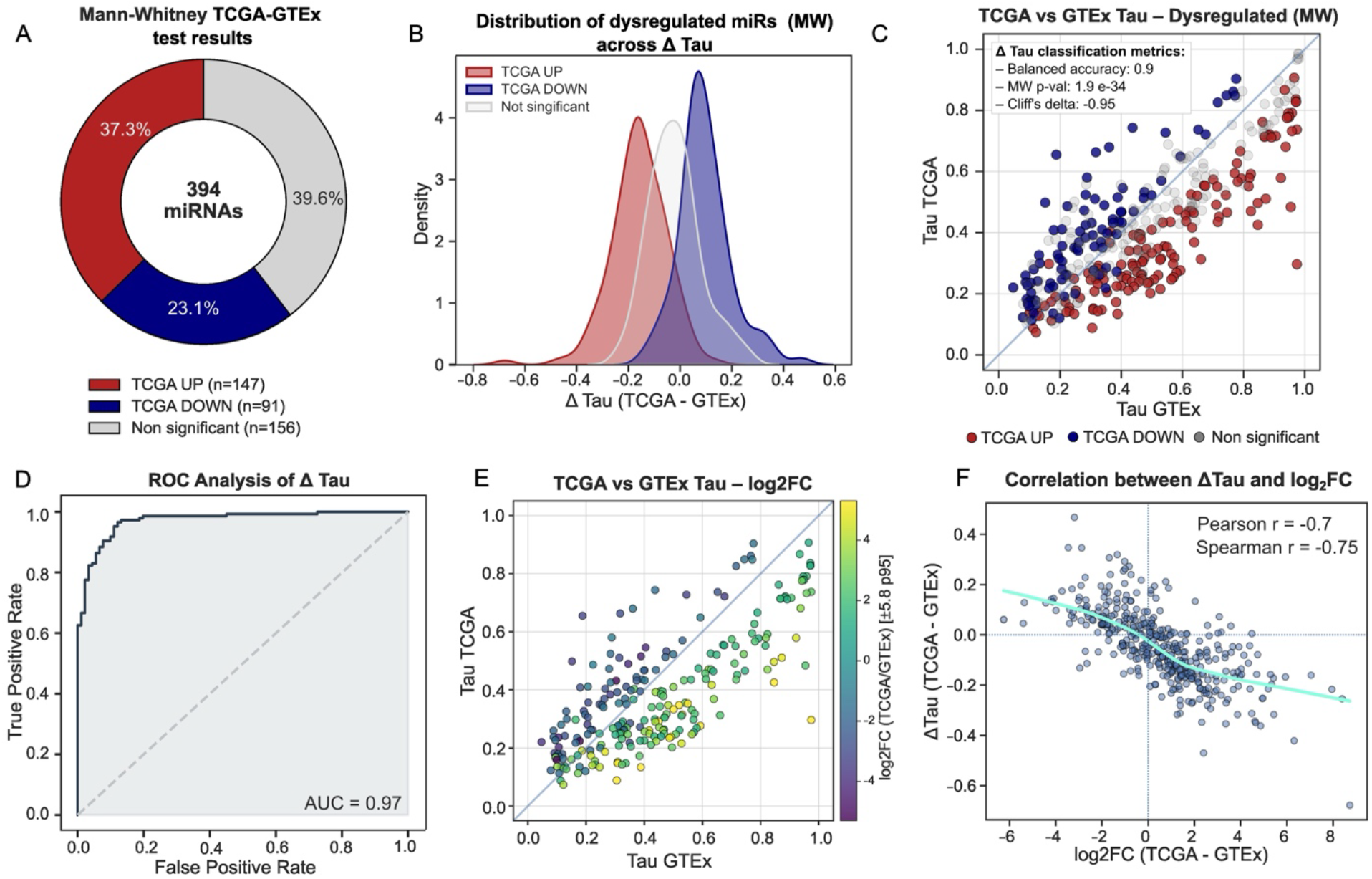
Association between cancer-associated expression shifts and tissue specificity dynamics. Distribution of miRNAs based on differential CPM expression between GTEx and TCGA cohorts across tissue types (Mann–Whitney FDR < 0.01; |log_2_FC| > 1). Among 394 analyzed miRNAs, 147 (37.3%) showed increased expression in cancer tissues (TCGA-upregulated), 91 (23.1%) showed increased expression in GTEx tissues (GTEx-upregulated), and 156 (39.6%) did not demonstrate significant expression differences. (B) Distribution of ΔTau values among the three miRNA classes (TCGA-upregulated, GTEx-upregulated, and non-significant miRNAs). Compared with recurrent dysregulation analysis, expression-based classification demonstrated a stronger separation of ΔTau distributions, with cancer-upregulated miRNAs preferentially showing reduced tissue specificity. (C) Scatter plot comparing tissue specificity scores (Tau) between GTEx and cancer cohorts. Each point represents an individual miRNA and is colored according to its expression-based differential expression status (TCGA-upregulated, GTEx-upregulated, or non-significant). The diagonal represents equal tissue specificity between normal and cancer cohorts; miRNAs below the diagonal exhibit reduced tissue specificity in cancer (negative ΔTau), whereas miRNAs above the diagonal exhibit increased tissue specificity (positive ΔTau). ΔTau showed strong discrimination between TCGA-upregulated and GTEx-upregulated miRNAs (balanced accuracy = 0.9; Mann–Whitney U test, P = 1.9 × 10^-34^; Cliff’s δ = −0.95). (D) Receiver operating characteristic (ROC) curve evaluating the ability of ΔTau to distinguish between TCGA-upregulated and GTEx-upregulated miRNAs. ΔTau demonstrated strong discriminatory performance between the two classes (ROC-AUC = 0.97). (E) Scatter plot comparing Tau scores between GTEx and cancer cohorts, with individual miRNAs colored according to log_2_ fold-change values. miRNAs with higher cancer-associated expression changes preferentially occupy the region corresponding to reduced tissue specificity (bottom right corner), whereas miRNAs with lower or negative expression changes tend toward preserved or increased tissue specificity (top left corner). (F) Correlation analysis between ΔTau and log_2_ fold-change values demonstrates a strong association between changes in tissue specificity and expression alterations (Pearson r = 0.7; Spearman ρ = 0.75).

### 3.3. Loss of tissue specificity and dysregulation

To identify the most robustly dysregulated miRNAs, we intersected the results of two independent analyses: (i) the directed binomial test summarizing differential expression across 17 TCGA cancer projects with matched normal tissues and (ii) the Mann–Whitney U test comparing mean miRNA expression between the GTEx and TCGA cohorts. Among the 394 analyzed miRNAs, 30 (7.6%) were consistently upregulated, 12 (3%) were consistently downregulated, whereas the remaining 352 (89.3%) did not satisfy both statistical criteria (Fig. 4A, B; Supplementary Fig. S2). From these robustly dysregulated miRNAs, we identified biologically relevant subset: normal-tissue-specific miRNAs that were consistently upregulated in cancer (n = 6; miR-519a-5p, miR-512-3p, miR-522-3p, miR-105-5p, miR-935, miR-1269a) (Fig. 4C).

**Fig 4.**
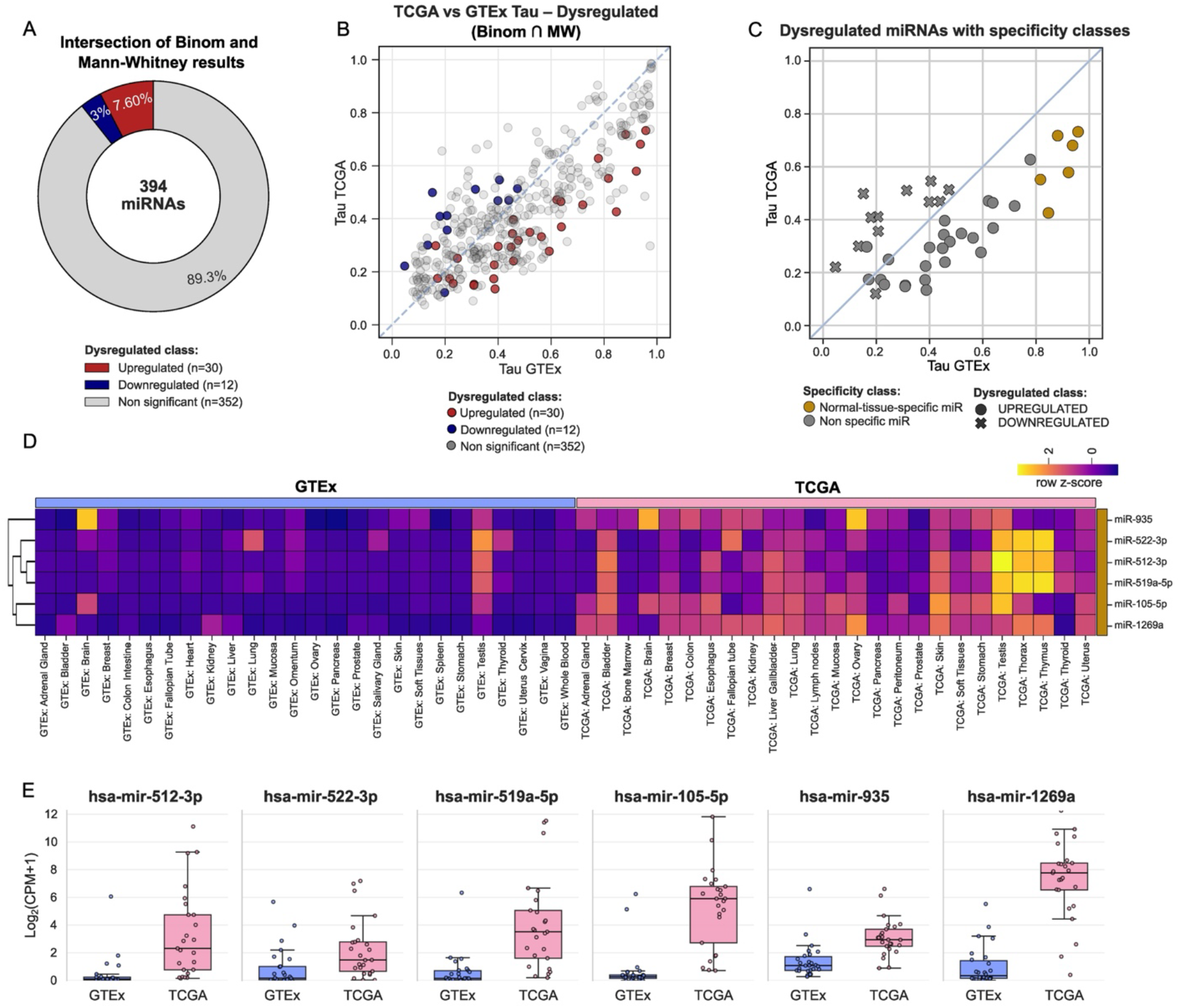
Integration of differential expression analyses and identification of miRNAs exhibiting altered tissue specificity during malignant transformation. (A) Distribution of miRNAs based on the intersection of two independent analyses: the directed binomial test across 17 TCGA cancer projects with matched normal tissues and comparison of mean CPM expression between GTEx and TCGA cohorts across tissue types. Among 394 analyzed miRNAs, 30 (7.6%) are constant upregulated, 30 (3%) are constant downregulated, and 352 (89.3%) are not significant. (B) Scatter plot comparing tissue specificity scores (Tau) between GTEx and TCGA cohorts. Each point represents an individual miRNA and is colored according to its dysregulation status based on the intersection of both statistical criteria. (C) Scatter plot highlighting dysregulated miRNAs according to their tissue specificity class. Yellow points indicate miRNAs exhibiting loss of tissue specificity during malignant transformation combined with consistent upregulation in cancer tissues, representing the primary candidates of interest. The purple cross indicates the miRNA showing acquired tissue specificity accompanied by consistent downregulation in cancer tissues. (D) Heatmap illustrating expression patterns of selected miRNAs with loss or acquired tissue specificity across GTEx and TCGA cohorts, demonstrating the transition of tissue expression patterns during malignant transformation. (E) Boxplots comparing log_2_ (CPM + 1) expression levels of selected miRNAs between GTEx and TCGA cohorts, illustrating differences in expression patterns associated with altered tissue specificity (Fig. 4D, E).

To further investigate the functional relevance of the identified miRNAs, pathway enrichment analysis was performed separately for each miRNA using experimentally validated target genes.

The numbers of validated targets varied substantially among the analyzed miRNAs: 54 for miR-519a-5p, 1,711 for miR-512-3p, 102 for miR-522-3p, 94 for miR-105-5p, 1,132 for miR-935, and 31 for miR-1269a. Significant pathway enrichment was detected only for miR-512-3p, miR-522-3p, miR-105-5p, and miR-935. The validated targets of miR-512-3p were significantly enriched in pathways associated with immune system functions and negative regulation of PI3K/AKT signaling. In contrast, miR-522-3p target genes did not show significant enrichment in metabolic or general biological pathways, the only significant associations were observed with the KEGG Prostate Cancer and Glioma pathways. Targets of miR-105-5p demonstrated enrichment in multiple cancer-related signaling pathways, including PI3K/AKT, EGFR, FoxO, mTOR, and AMPK signaling, as well as in pathways involved in the regulation of stem cell pluripotency. The target genes of miR-935 were significantly associated with a broad range of biological processes and signaling pathways, including response to stress, negative regulation of gene expression, response to cytokines, blood vessel development, cell death, programmed cell death, apoptotic processes, transcriptional regulation by TP53, immune system, and signaling by NTRKs (Supplementary Table S3).

## 4. Discussion

This study presents an integrative analysis of microRNA (miRNA) biology by combining tissue specificity with differential expression across healthy and malignant tissues. This approach characterizes miRNAs not only by changes in expression but also by alterations in tissue-specific regulatory programs during malignant transformation, a dimension that has received relatively little attention in previous pan-cancer studies. Tissue specificity was quantified using the Tau index, a robust and widely used metric of expression specificity. To our knowledge, this is the first large-scale study to systematically examine the relationship between miRNA dysregulation and changes in tissue specificity across healthy and cancerous tissues.

In contrast to previous pan-cancer studies that primarily relied on differential expression analysis between tumor and matched normal samples to identify potential oncogenic or tumor-suppressive miRNAs [13-15], we incorporated normal tissue expression (GTEx) as an independent validation layer. This approach was implemented to minimize false-positive findings that may arise from recurrence-based analyses alone. A representative example is miR-671-5p. Based solely on recurrence across TCGA projects, this miRNA appeared to function as a consistent oncomiR, being upregulated in 11 of 17 cancer types and downregulated in none. However, direct comparison of global mean expression demonstrated significantly higher expression in normal tissues (GTEx) than in tumors (TCGA), consistent with previous studies describing miR-671-5p primarily as a tumor suppressor in multiple malignancies [16-20]. Similar discrepancies were observed for six additional miRNAs, including miR-122-5p, miR-9-3p, miR-760, miR-148b-5p, and miR-135b-5p. Conversely, three miRNAs (miR-29c-3p, miR-30a-5p, and miR-140-3p) were classified as recurrently downregulated despite exhibiting significantly higher global mean expression in the cancer cohort. Applying concordance between recurrence-based and expression-based analyses excluded these contradictory cases and resulted in a more conservative but higher-confidence set of dysregulated miRNAs. Similar borderline classifications have been reported in previous pan-cancer analyses. For example, Sharif Moradi et al. identified miR-206 and miR-381 as normomiRs with potential tumor-suppressive properties. In our analysis, miR-206 demonstrated recurrent downregulation (7 downregulated versus 1 upregulated, binom p-val = 0.04) but did not show a statistically significant difference between GTEx and TCGA global expression levels. Conversely, miR-381-3p exhibited significantly higher expression in GTEx tissues compared with TCGA, but its recurrence pattern (7 downregulated versus 2 upregulated, binom p-val = 0.09) did not reach statistical significance according to the binomial test. Therefore, we used a two-step validation approach requiring agreement between independent dysregulation criteria to define a high-confidence set of cancer-associated miRNAs.

Our results suggest that changes in tissue specificity are associated, at least in part, with miRNA dysregulation. This observation is biologically plausible, as miRNAs involved in tissue-specific functions may acquire more widespread activity during malignant transformation and contribute to common oncogenic processes. From a therapeutic perspective, such miRNAs may represent attractive candidates for anti-miRNA strategies, since their limited expression in normal tissues could reduce off-target effects associated with systemic inhibition. We identified six miRNA that loss tissue specificity and displayed robust upregulation pattern in cancer. The functional relevance of this subset is supported by published experimental evidence. Six miRNAs within this signature have documented oncogenic activity across multiple malignancies. MiR-522-3p promotes proliferation in osteosarcoma [21] and enhances tumor growth, metastasis, and epithelial-mesenchymal transition (EMT) in colorectal cancer [22, 23]. Although evidence for miR-519a-5p remains limited, it has been reported to increase the migratory and invasive capacity of hepatocellular carcinoma cells [24], whereas an opposite effect, namely inhibition of the Warburg phenotype, has been described in ovarian cancer [25]. MiR-105-5p has also been consistently implicated in tumor progression by promoting EMT and immunosuppression in bladder [26], breast [27], and gastric cancers [28]. Similarly, miR-1269a exhibits predominantly oncogenic activity, promoting EMT in colorectal and bladder cancers [29, 30], enhancing migration and proliferation in esophageal squamous cell carcinoma [31], and stimulating glioma progression while suppressing apoptosis in vitro [32]. In contrast, miR-512-3p and miR-935 display context-dependent functions. MiR-512-3p has been associated with poor clinical outcomes in pediatric medulloblastoma [33] and shown to activate the PI3K/AKT and Wnt/β-catenin signaling pathways in colorectal cancer [34]. It has also been implicated in the regulation of chemoresistance in ovarian cancer through the miR-512-3p/RPS6KA2 axis. Conversely, other studies reported that the miR-512-3p/HK-2 axis suppresses tumor progression in ovarian [35], prostate [36], and colorectal cancers [37]. Likewise, miR-935 has been reported to stimulate proliferation in hepatocellular and gastric cancers and to promote chemoresistance in lung cancer through the miR-935/SOX7 axis [38-40]. Similar oncogenic effects have also been described in choriocarcinoma via the miR-935/GJA1 axis [41]. In contrast, the miR-935/HIF1α axis has been shown to suppress glioma progression [42], and miR-935 inhibits proliferation, migration, and invasion in oral squamous cell carcinoma [43].

Although clinical experience with miRNA inhibitors (antagomirs/anti-miRs) in oncology remains limited to early-phase trials, available evidence indicates a generally favorable safety profile. Cobomarsen (MRG-106), an LNA inhibitor of miR-155, was well tolerated in Phase 1, while the subsequent Phase 2 SOLAR trial was terminated for organizational reasons unrelated to safety or efficacy (NCT02580552, NCT03713320). Likewise, the first-in-class LNA-i-miR-221 demonstrated no grade 3–4 toxicities and preliminary antitumor activity in a first-in-human Phase 1 study [44], and the antisense miR-10b inhibitor TTX-MC138 has not shown dose-limiting toxicities in an ongoing Phase 1/2 trial (NCT06260774). In contrast, clinical development of the miR-34a mimic MRX34 was discontinued because of severe immune-mediated toxicity [45, 46]. Notably, both miR-155-5p and miR-10b exhibit low tissue specificity and are broadly expressed in healthy tissues, yet their pharmacological inhibition has remained clinically tolerable. Together, these findings support the feasibility of targeting oncogenic miRNAs and suggest that candidates with higher baseline tissue specificity, such as those identified in this study, may offer an even wider therapeutic window. Notably, miR-155-5p and both miR-10b isoforms, which have already entered clinical development as therapeutic targets, exhibit relatively low tissue specificity and are broadly expressed in healthy tissues. Despite this, inhibition of these miRNAs has shown acceptable tolerability in clinical studies. In comparison, the candidates identified in our study combine broad upregulation across multiple malignancies with substantially higher tissue specificity under normal physiological conditions. Although this hypothesis requires experimental validation, these characteristics suggest that their therapeutic inhibition could provide an even wider safety margin while targeting molecular pathways shared across multiple cancer types.

Several limitations should be considered when interpreting these findings. First, the tissue compositions of the TCGA and GTEx cohorts differ, despite having comparable numbers of tissue types (25 and 26, respectively). For instance, TCGA includes tissues such as bone marrow, lymph nodes, peritoneum, thorax, and thymus, which are not represented in GTEx. Conversely, GTEx contains tissues including the heart, omentum, salivary gland, spleen, vagina, and whole blood, which are absent from TCGA. These differences in tissue representation may affect the assessment of tissue specificity and potentially influence the observed results. Additionally, the sizes of the selected miRNA target sets (n = 6) used for functional analysis varied substantially. For example, 1,711 targets meeting the inclusion criteria were identified for miR-512-3p, whereas only 31 targets were identified for miR-1269a. This considerable variation in target set size likely contributed to the limited success of functional enrichment analyses, which yielded sufficient results only for three miRNAs: miR-512-3p, miR-105-5p, and miR-935.

## 5. Declaration of generative AI and AI-assisted technologies in the writing process

During the preparation of this manuscript, the authors used ChatGPT (OpenAI) solely for the purpose of improving language quality, grammar, and readability. The authors carefully reviewed and edited all AI-assisted suggestions and take full responsibility for the content of the published article.

## 6. Acknowledgements

## 7. Author contributions

AI: Conceptualization, Data curation, Formal analysis, Investigation, Methodology, Visualization, Writing – original draft.

MP: Conceptualization, Funding acquisition, Methodology, Project administration, Supervision, Writing – review & editing.

AB: Funding acquisition, Writing – review & editing.

AE: Data curation.

All authors read and approved the final manuscript.

## 8. Data and code availability

The datasets used in this study are publicly available. MicroRNA expression data were obtained from The Cancer Genome Atlas (TCGA) and the Genotype-Tissue Expression (GTEx) project. These datasets can be accessed through their respective public repositories. The code used for data processing, analysis, and generation of the results presented in this study is available at: https://github.com/neuropromotion/Bioinformatics/tree/main/miRNA_bulk.

