## Supplementary Figure 1 for "A pan-cancer analysis of microRNA tissue specificity and its association with dysregulation"

**Fig S1. Volcano plot analysis of differential expression between tumor and normal tissues across TCGA, TARGET, CGCI, and CPTAC cohorts.**

The figure presents 17 volcano plots summarizing miRNA differential expression between tumor and corresponding normal tissues across multiple cancer types. The analysis spans a broad spectrum of malignancies, including acute myeloid leukemia (TARGET-AML), breast carcinoma (TCGA-BRCA), breast cancer proteogenomic cohort (CPTAC-3), endometrial carcinoma (TCGA-UCEC), clear cell renal cell carcinoma (TCGA-KIRC), lung adenocarcinoma (TCGA-LUAD), head and neck squamous cell carcinoma (TCGA-HNSC), thyroid carcinoma (TCGA-THCA), prostate adenocarcinoma (TCGA-PRAD), lung squamous cell carcinoma (TCGA-LUSC), gastric adenocarcinoma (TCGA-STAD), bladder urothelial carcinoma (TCGA-BLCA), hepatocellular carcinoma (TCGA-LIHC), papillary and chromophobe renal cell carcinomas (TCGA-KIRP, TCGA-KICH), esophageal carcinoma (TCGA-ESCA), Burkitt lymphoma (CGCI-BLGSP). Each plot illustrates the global pattern of up- and downregulated miRNAs distinguishing tumor from normal tissue within each cancer type.

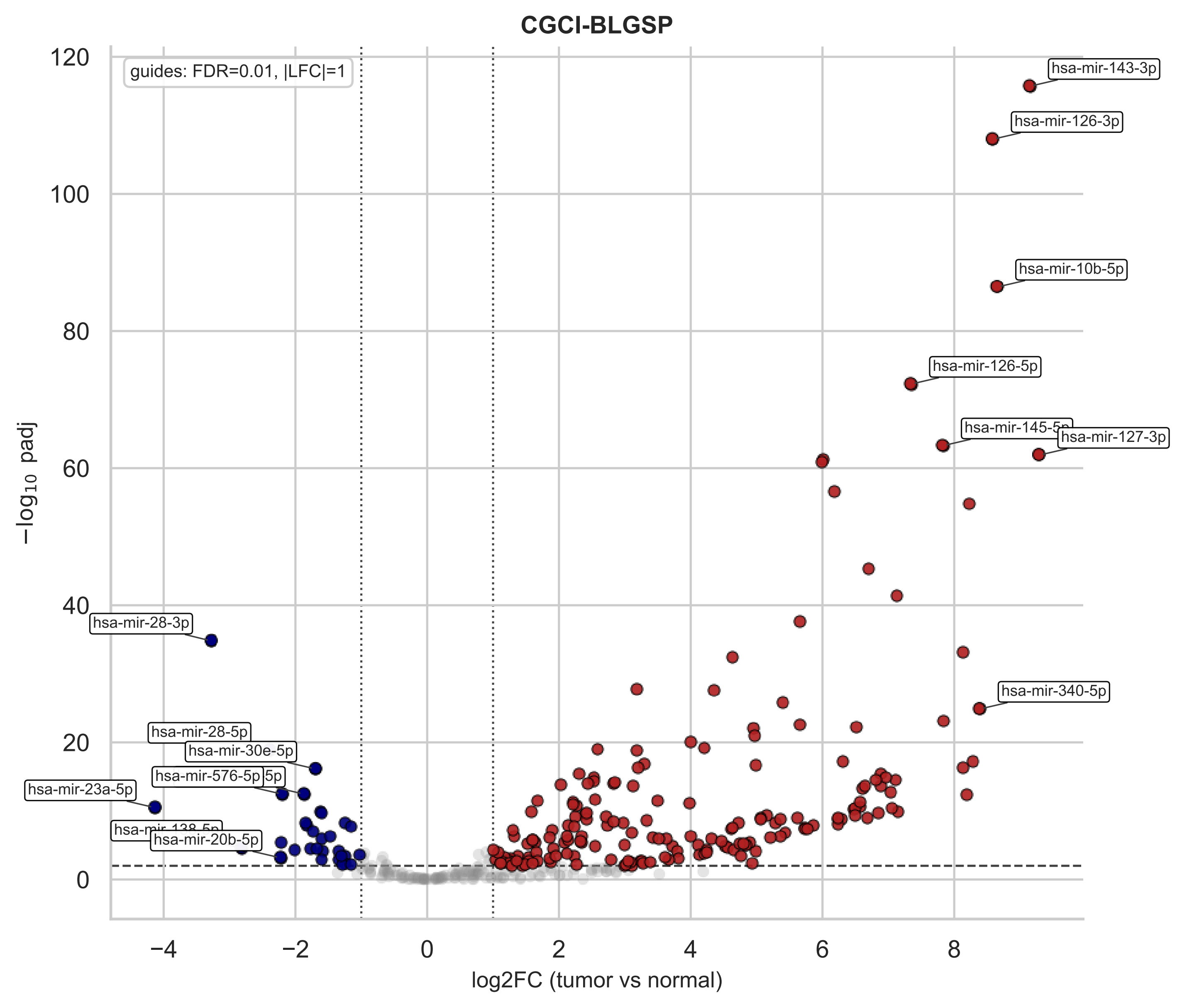

CGCI-BLGSP
guides: FDR=0.01, |LFC|=1
hsa-mir-143-3p
hsa-mir-126-3p
hsa-mir-10b-5p
hsa-mir-126-5p
hsa-mir-127-3p
hsa-mir-340-5p
hsa-mir-28-3p
hsa-mir-28-5p
hsa-mir-30e-5p
hsa-mir-576-5p
hsa-mir-23a-5p
hsa-mir-20b-5p
−log10 padj
log2FC (tumor vs normal)

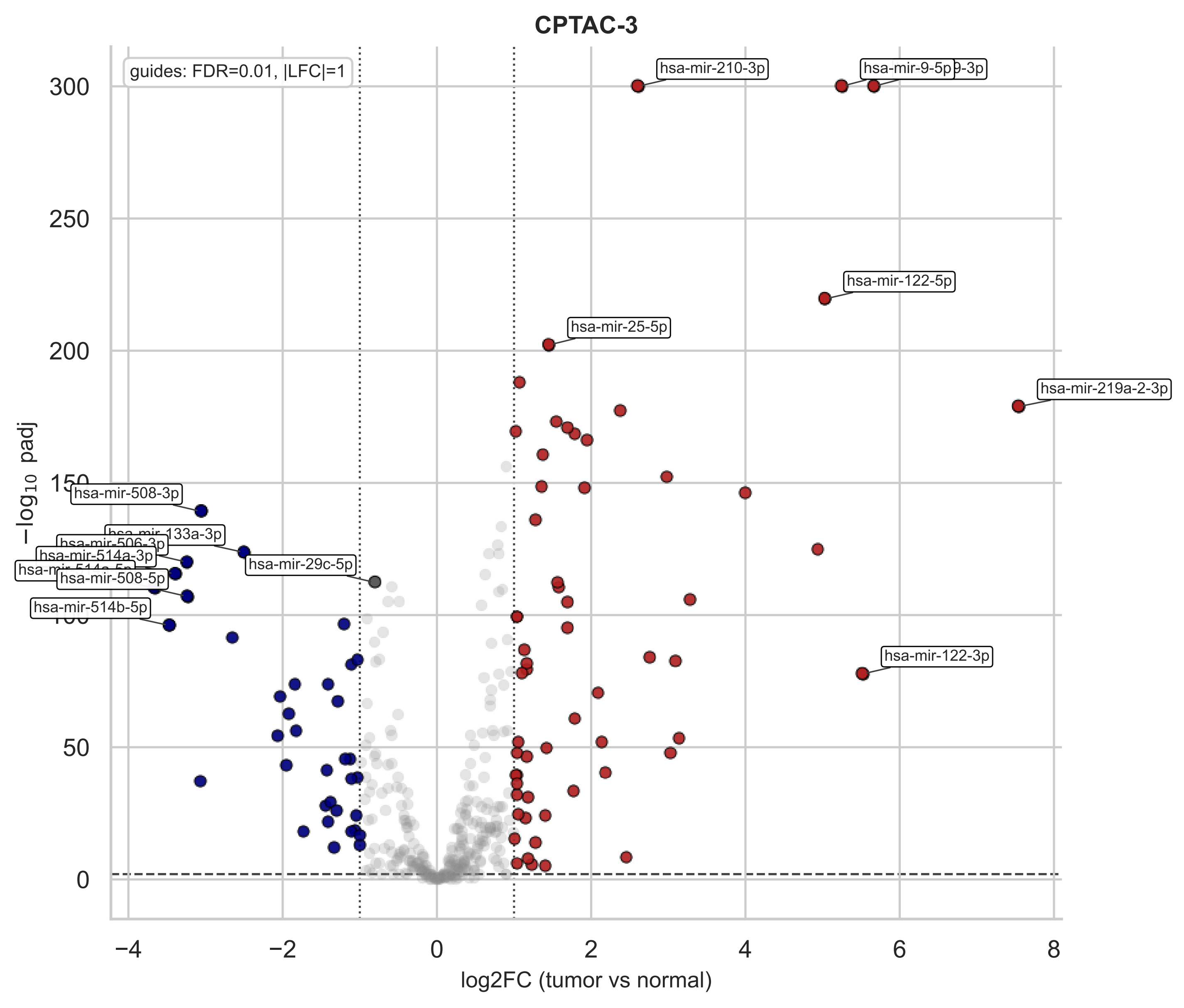
CPTAC-3
guides: FDR=0.01, |LFC|=1
hsa-mir-210-3p
hsa-mir-9-5p
hsa-mir-122-5p
hsa-mir-25-5p
hsa-mir-219a-2-3p
hsa-mir-508-3p
hsa-mir-133a-3p
hsa-mir-506-3p
hsa-mir-514a-3p
hsa-mir-508-5p
hsa-mir-29c-5p
hsa-mir-514b-5p
hsa-mir-122-3p
300
250
200
150
100
50
0
−4
−2
0
2
4
6
8
log2FC (tumor vs normal)
−log10 padj

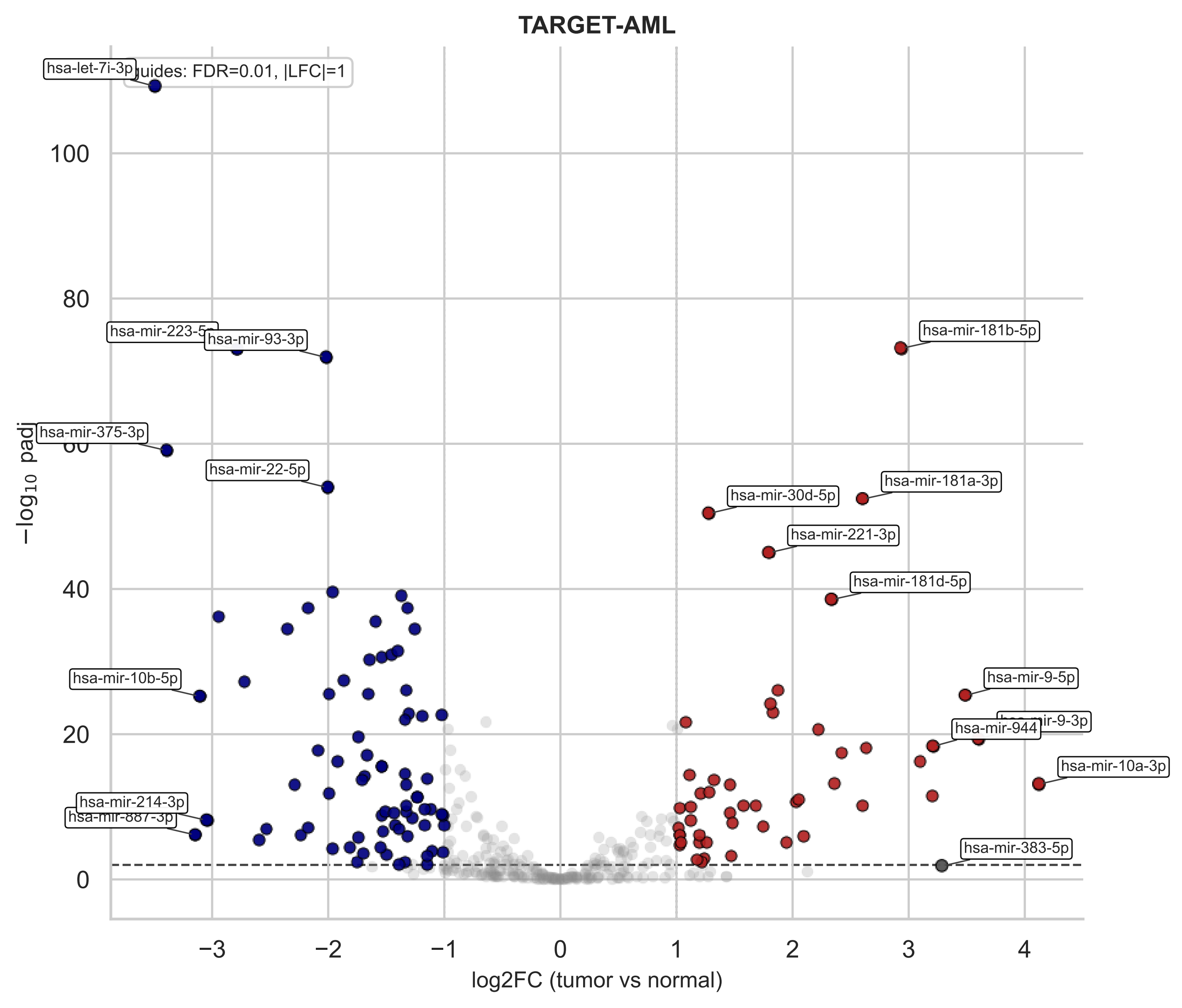

TARGET-AML
guides: FDR=0.01, |LFC|=1
hsa-let-7i-3p
hsa-mir-93-3p
hsa-mir-375-3p
hsa-mir-22-5p
hsa-mir-10b-5p
hsa-mir-214-3p
hsa-mir-181b-5p
hsa-mir-181a-3p
hsa-mir-30d-5p
hsa-mir-221-3p
hsa-mir-181d-5p
hsa-mir-9-5p
hsa-mir-944
hsa-mir-10a-3p
hsa-mir-383-5p
−log10 padj
log2FC (tumor vs normal)

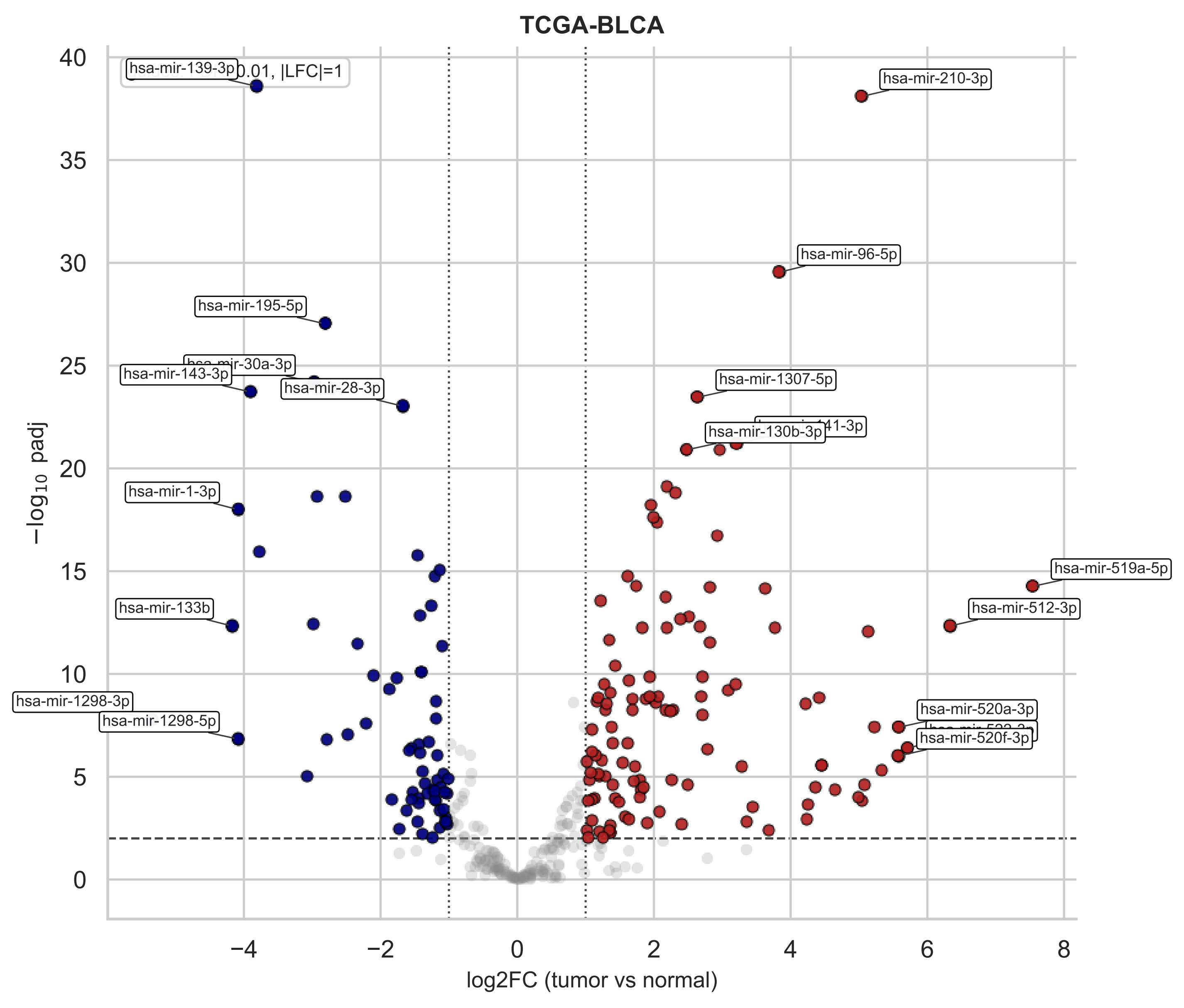
TCGA-BLCA
0.01, |LFC|=1
hsa-mir-139-3p
hsa-mir-195-5p
hsa-mir-143-3p
hsa-mir-28-3p
hsa-mir-1-3p
hsa-mir-133b
hsa-mir-1298-3p
hsa-mir-1298-5p
hsa-mir-210-3p
hsa-mir-96-5p
hsa-mir-1307-5p
hsa-mir-130b-3p
hsa-mir-519a-5p
hsa-mir-512-3p
hsa-mir-520a-3p
hsa-mir-520f-3p
−log10 padj
log2FC (tumor vs normal)
−4
−2
0
2
4
6
8
0
5
10
15
20
25
30
35
40

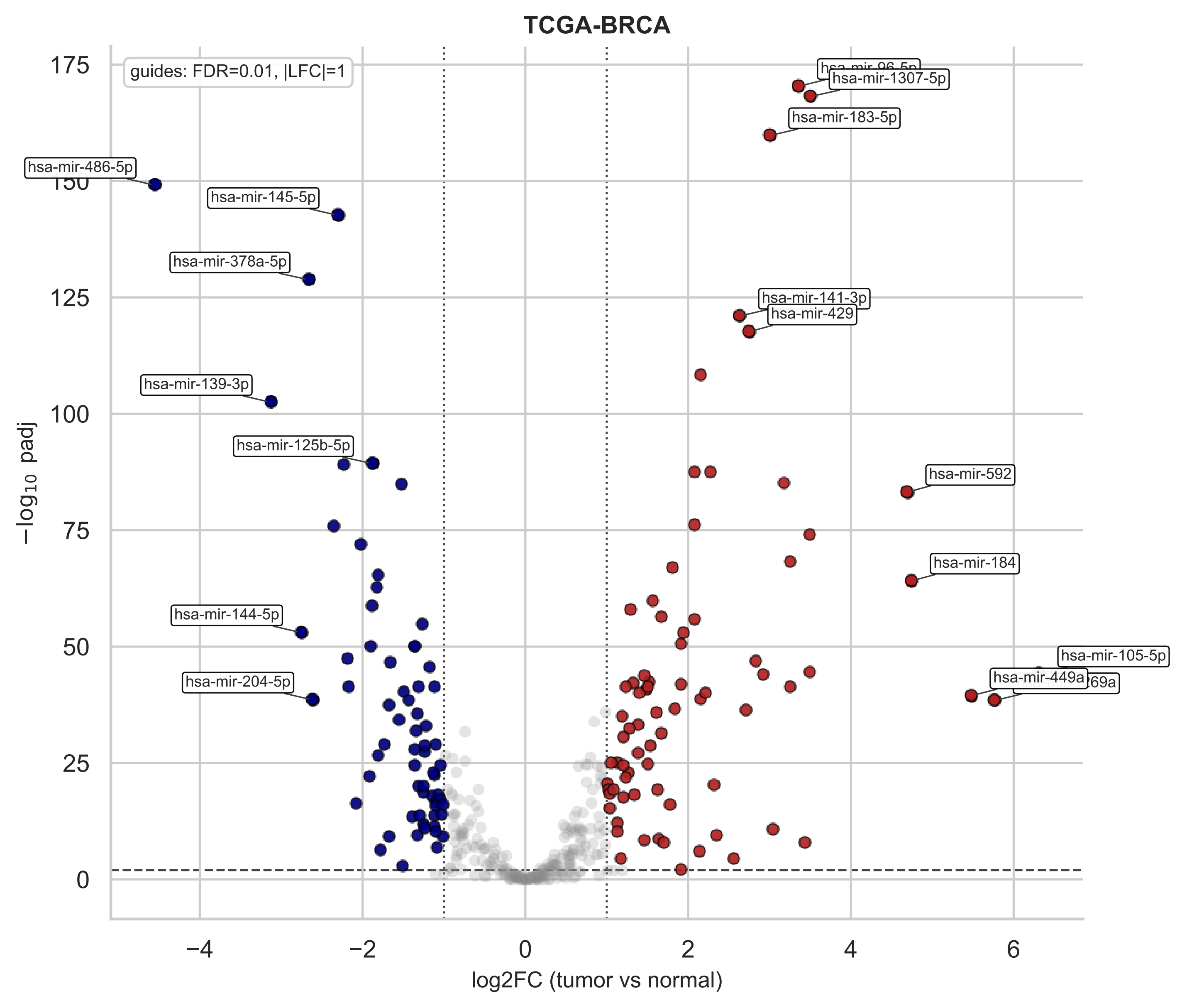
TCGA-BRCA
guides: FDR=0.01, |LFC|=1
hsa-mir-1307-5p
hsa-mir-183-5p
hsa-mir-486-5p
hsa-mir-145-5p
hsa-mir-378a-5p
hsa-mir-141-3p
hsa-mir-429
hsa-mir-139-3p
hsa-mir-125b-5p
hsa-mir-592
hsa-mir-184
hsa-mir-144-5p
hsa-mir-105-5p
hsa-mir-449a
hsa-mir-204-5p
175
150
125
100
75
50
25
0
$-\log_{10}$ padj
−4
−2
0
2
4
6
log2FC (tumor vs normal)

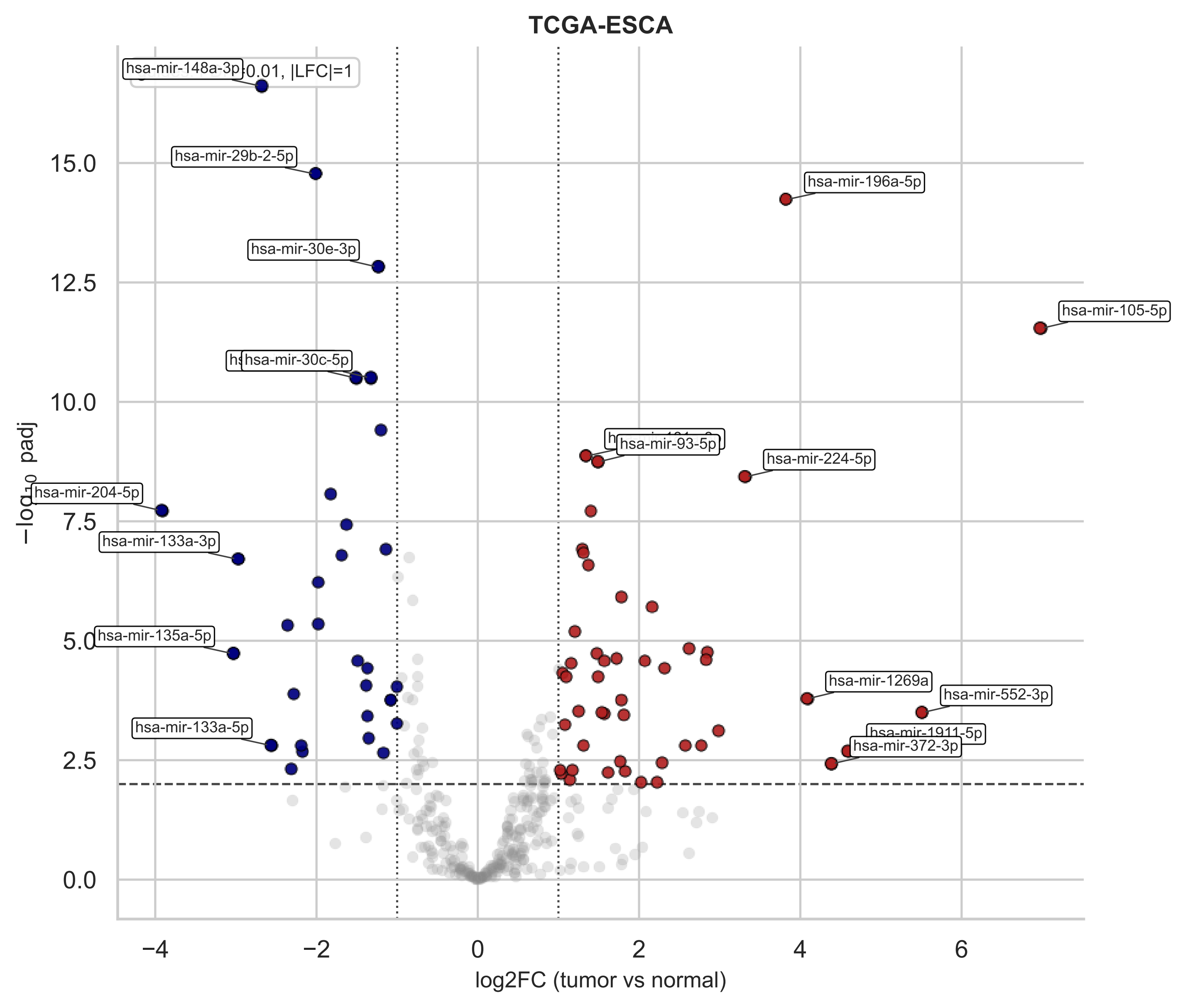

TCGA-ESCA
hsa-mir-148a-3p
0.01, |LFC|=1
hsa-mir-29b-2-5p
hsa-mir-196a-5p
hsa-mir-30e-3p
hsa-mir-105-5p
hsa-mir-30c-5p
hsa-mir-93-5p
hsa-mir-224-5p
hsa-mir-204-5p
hsa-mir-133a-3p
hsa-mir-135a-5p
hsa-mir-1269a
hsa-mir-552-3p
hsa-mir-133a-5p
hsa-mir-1911-5p
hsa-mir-372-3p
−log10 padj
log2FC (tumor vs normal)

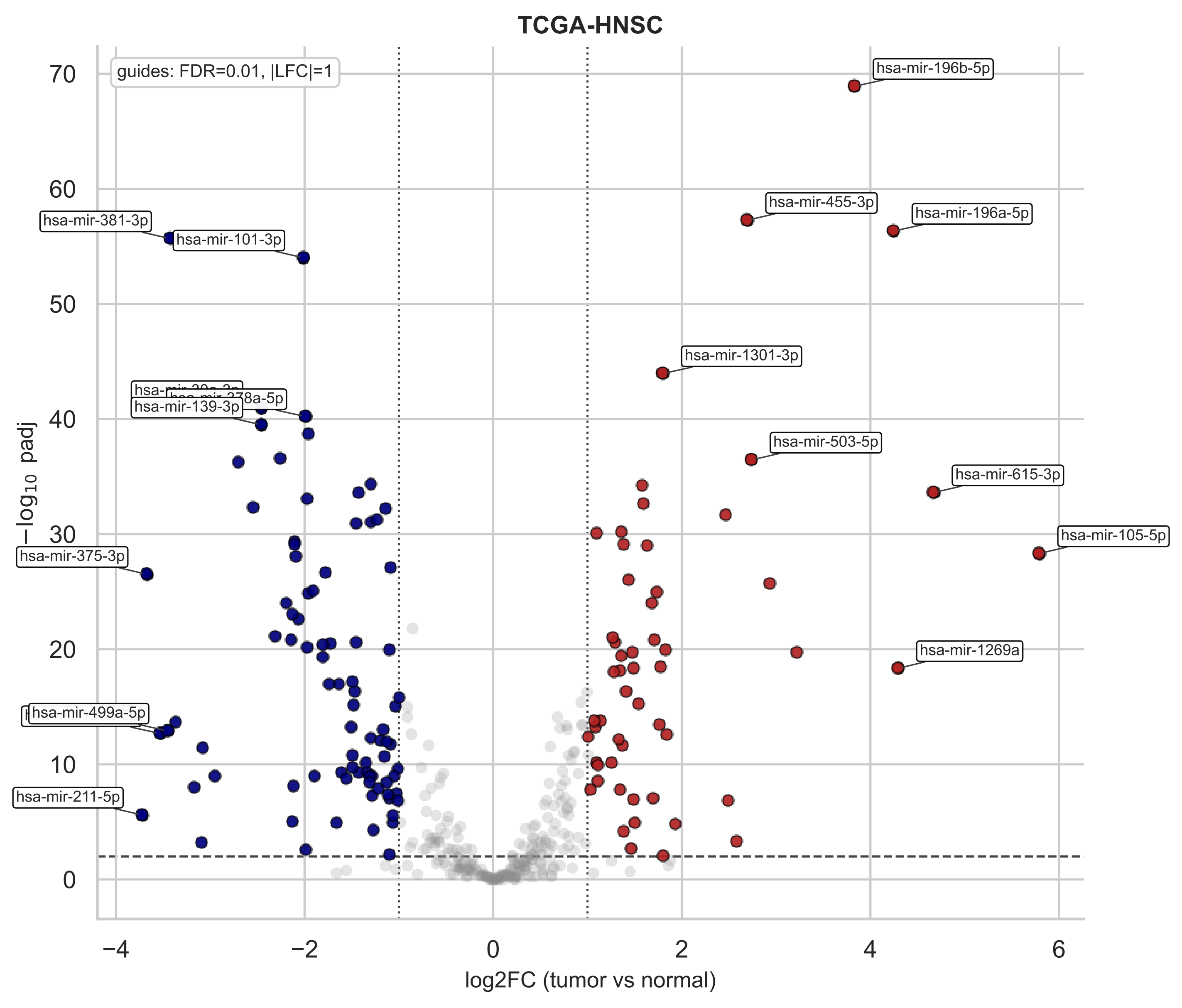
TCGA-HNSC
guides: FDR=0.01, |LFC|=1
−log10 padj
log2FC (tumor vs normal)
hsa-mir-196b-5p
hsa-mir-455-3p
hsa-mir-196a-5p
hsa-mir-381-3p
hsa-mir-101-3p
hsa-mir-1301-3p
hsa-mir-139-3p
hsa-mir-503-5p
hsa-mir-615-3p
hsa-mir-105-5p
hsa-mir-375-3p
hsa-mir-1269a
hsa-mir-499a-5p
hsa-mir-211-5p

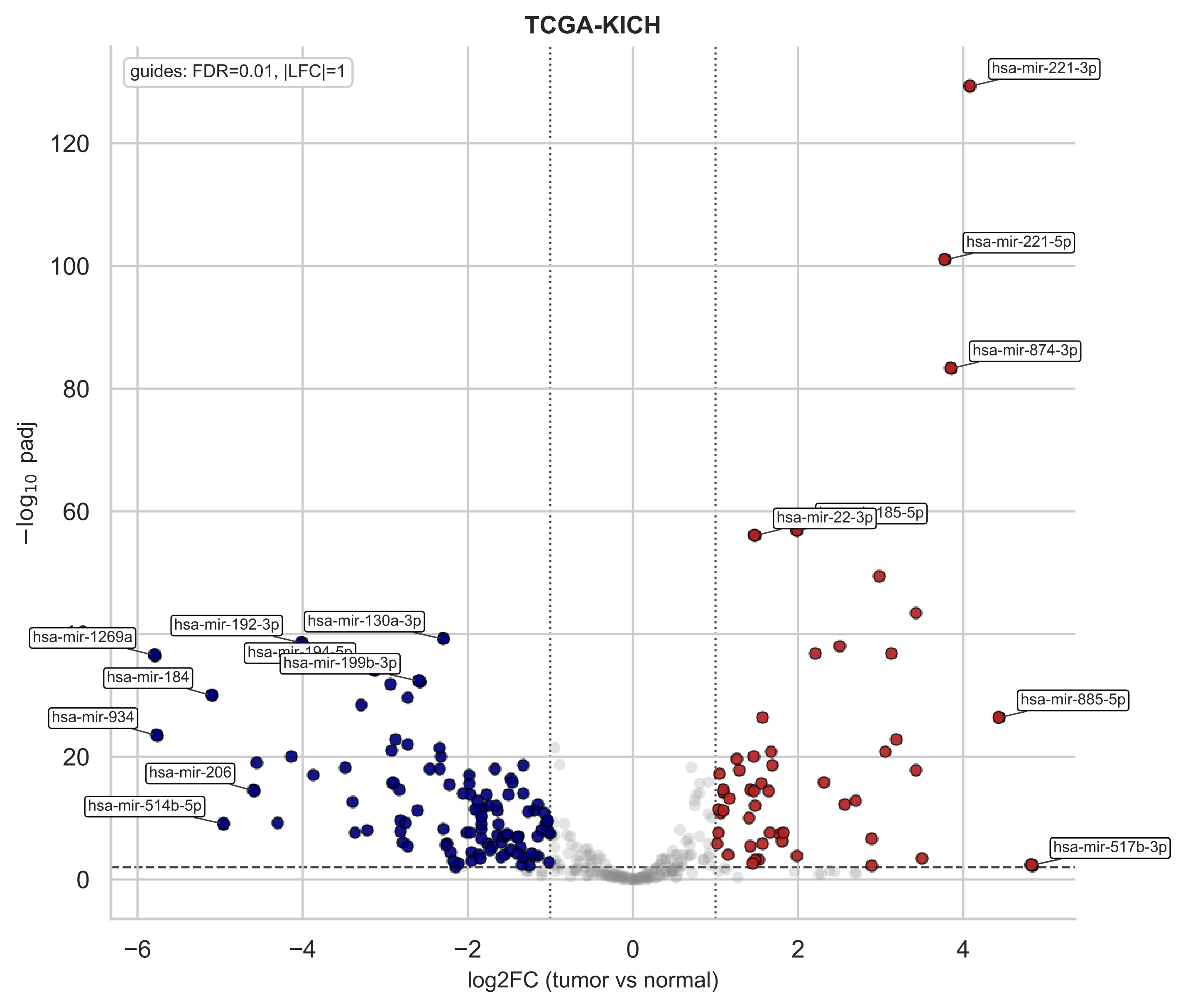
TCGA-KICH
guides: FDR=0.01, |LFC|=1
hsa-mir-221-3p
hsa-mir-221-5p
hsa-mir-874-3p
hsa-mir-22-3p
hsa-mir-885-5p
hsa-mir-517b-3p
hsa-mir-1269a
hsa-mir-192-3p
hsa-mir-130a-3p
hsa-mir-199b-3p
hsa-mir-184
hsa-mir-934
hsa-mir-206
hsa-mir-514b-5p
−log10 padj
log2FC (tumor vs normal)

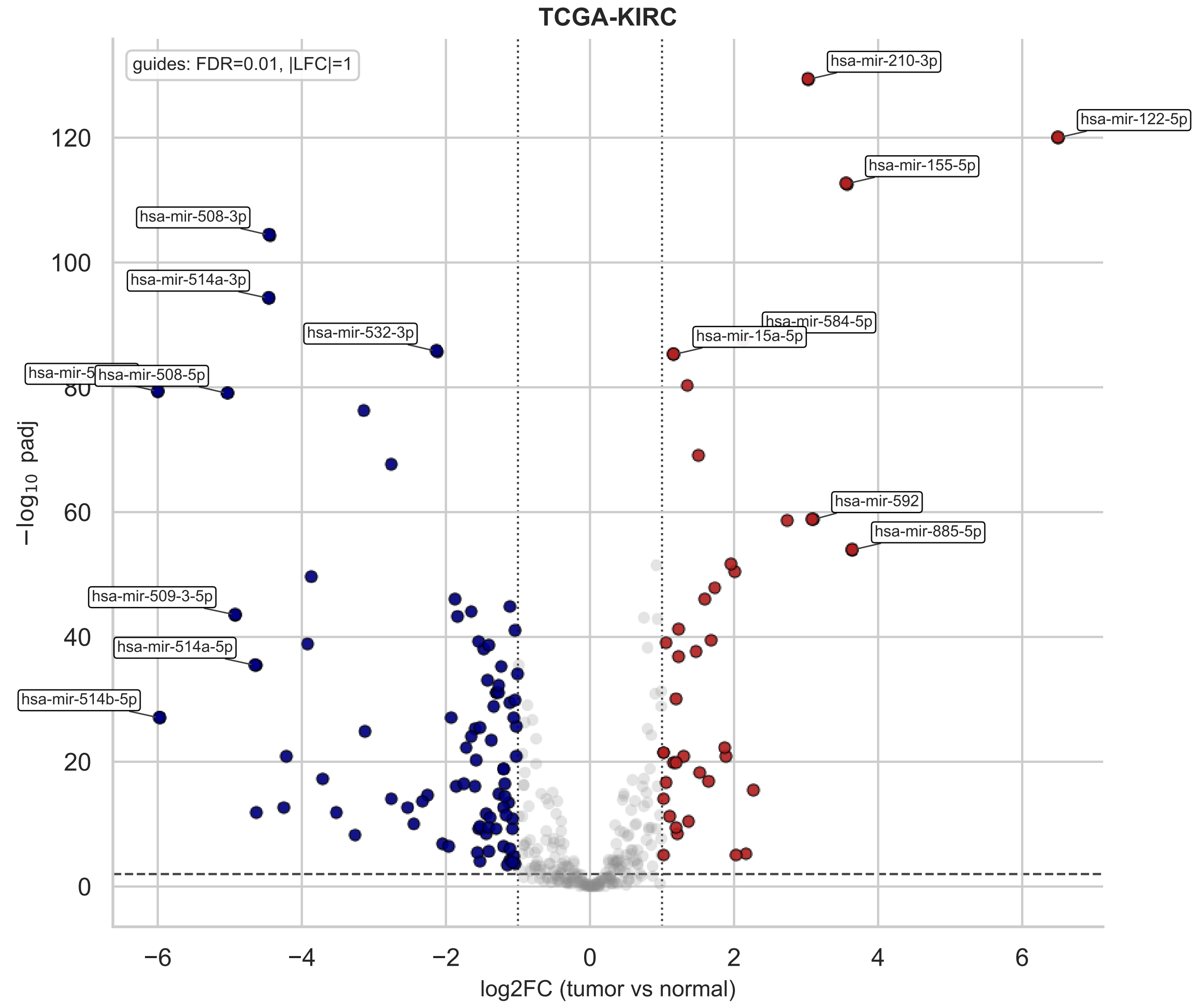
TCGA-KIRC
guides: FDR=0.01, |LFC|=1
hsa-mir-210-3p
hsa-mir-122-5p
hsa-mir-155-5p
hsa-mir-508-3p
hsa-mir-514a-3p
hsa-mir-532-3p
hsa-mir-584-5p
hsa-mir-15a-5p
hsa-mir-508-5p
hsa-mir-592
hsa-mir-885-5p
hsa-mir-509-3-5p
hsa-mir-514a-5p
hsa-mir-514b-5p
−log10 padj
log2FC (tumor vs normal)
0
20
40
60
80
100
120
−6
−4
−2
0
2
4
6

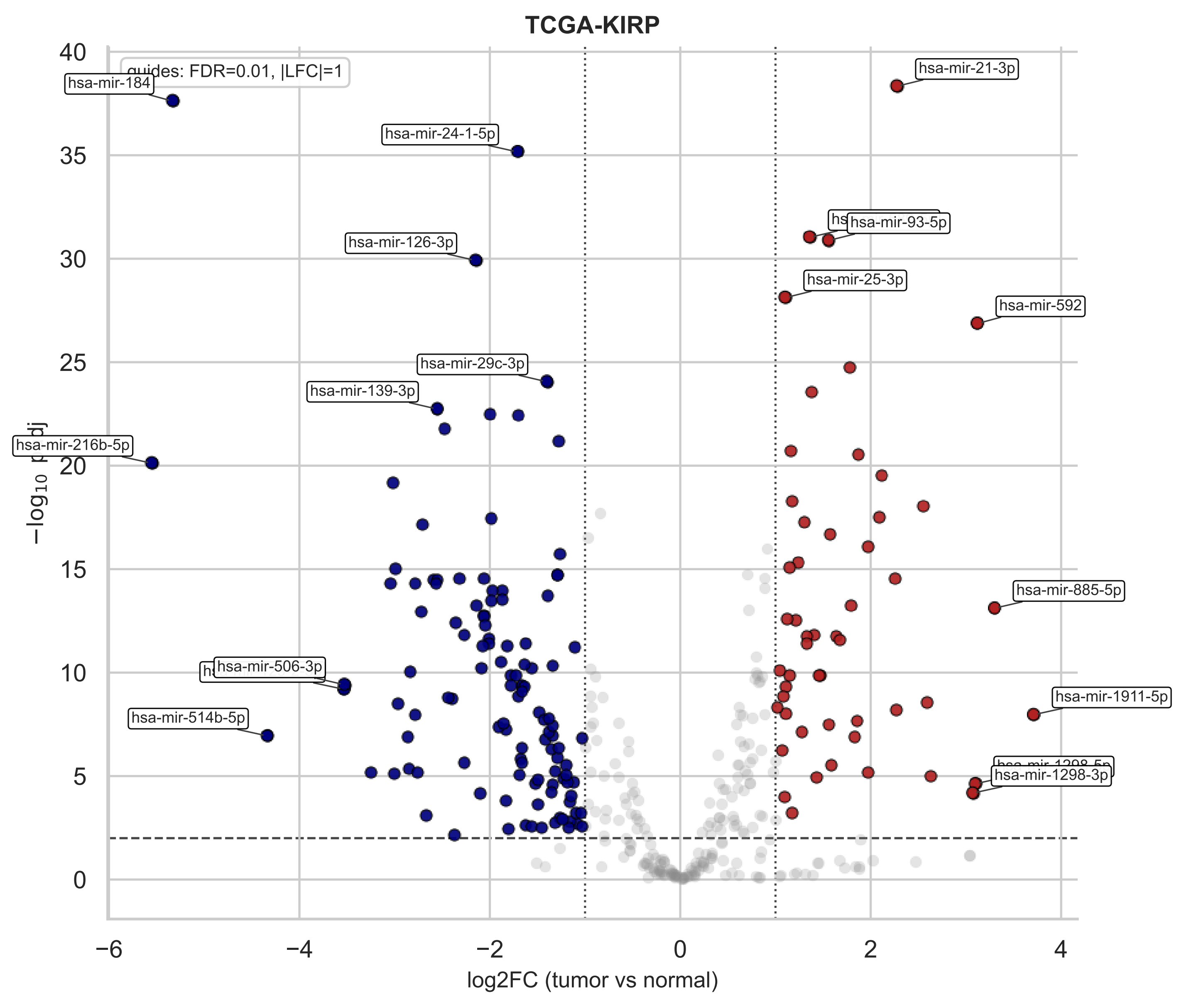

TCGA-KIRP
guides: FDR=0.01, |LFC|=1
hsa-mir-184
hsa-mir-24-1-5p
hsa-mir-126-3p
hsa-mir-29c-3p
hsa-mir-139-3p
hsa-mir-216b-5p
hsa-mir-506-3p
hsa-mir-514b-5p
hsa-mir-21-3p
hsa-mir-93-5p
hsa-mir-25-3p
hsa-mir-592
hsa-mir-885-5p
hsa-mir-1911-5p
hsa-mir-1298-3p
$-\log_{10}$ padj
log2FC (tumor vs normal)
40
35
30
25
20
15
10
5
0
−6
−4
−2
0
2
4

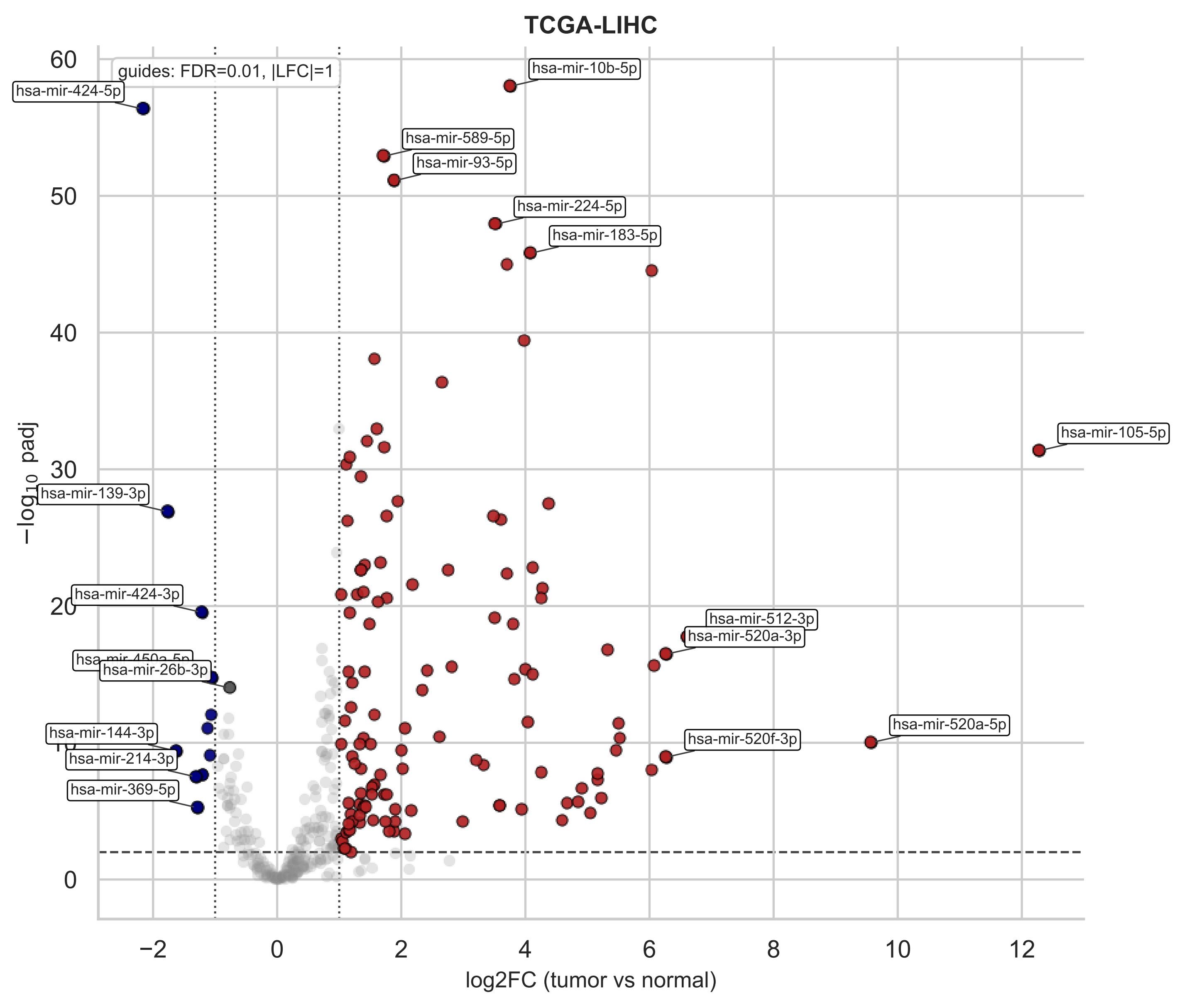

TCGA-LIHC
guides: FDR=0.01, |LFC|=1
hsa-mir-424-5p
hsa-mir-10b-5p
hsa-mir-589-5p
hsa-mir-93-5p
hsa-mir-224-5p
hsa-mir-183-5p
hsa-mir-105-5p
hsa-mir-139-3p
hsa-mir-424-3p
hsa-mir-512-3p
hsa-mir-520a-3p
hsa-mir-26b-3p
hsa-mir-144-3p
hsa-mir-214-3p
hsa-mir-369-5p
hsa-mir-520a-5p
hsa-mir-520f-3p
−log10 padj
log2FC (tumor vs normal)

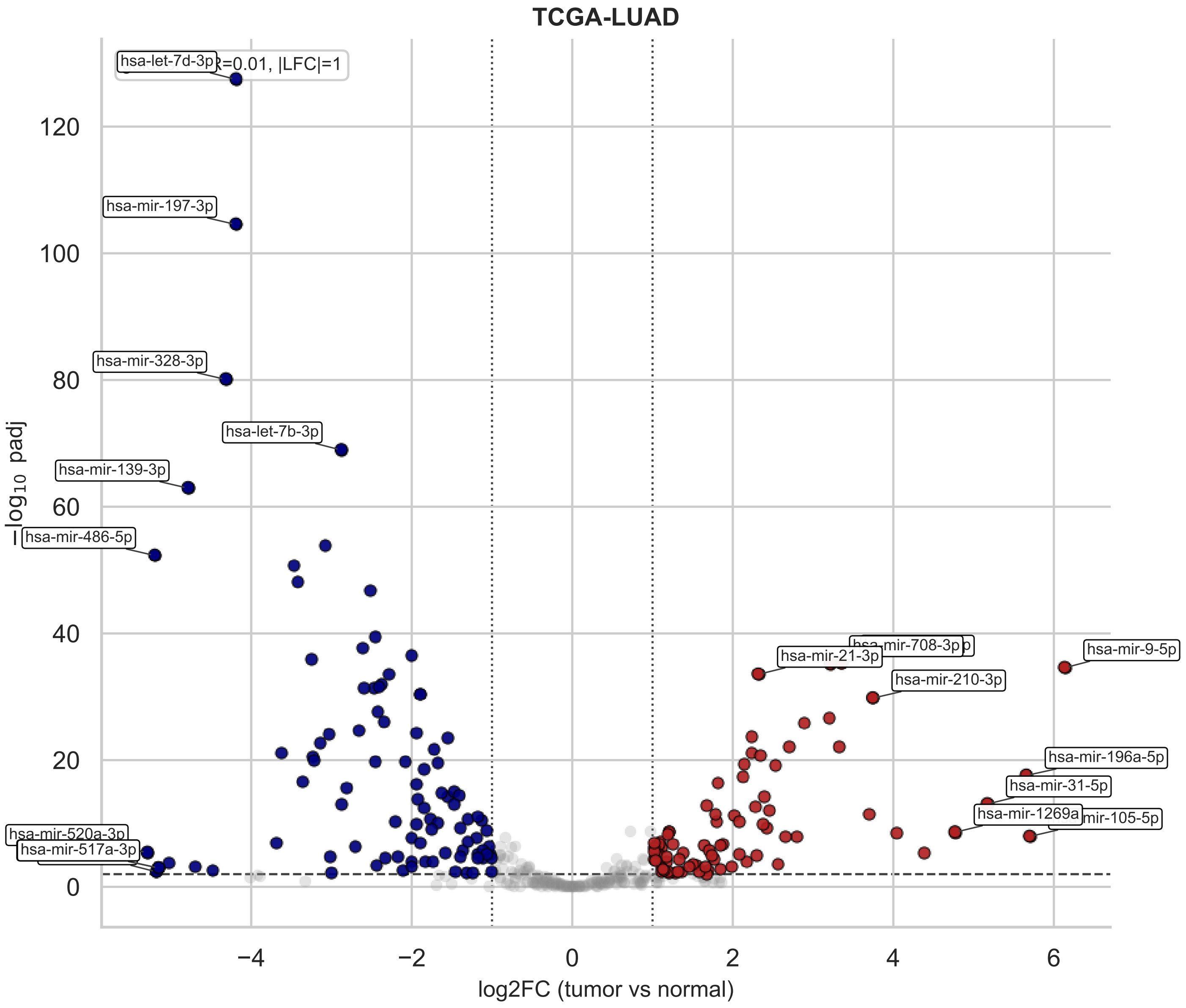
TCGA-LUAD
R=0.01, |LFC|=1
hsa-let-7d-3p
hsa-mir-197-3p
hsa-mir-328-3p
hsa-let-7b-3p
hsa-mir-139-3p
hsa-mir-486-5p
hsa-mir-520a-3p
hsa-mir-517a-3p
hsa-mir-708-3p
hsa-mir-21-3p
hsa-mir-9-5p
hsa-mir-210-3p
hsa-mir-196a-5p
hsa-mir-31-5p
hsa-mir-1269a
mir-105-5p
−log10 padj
log2FC (tumor vs normal)
120
100
80
60
40
20
0
−4
−2
0
2
4
6

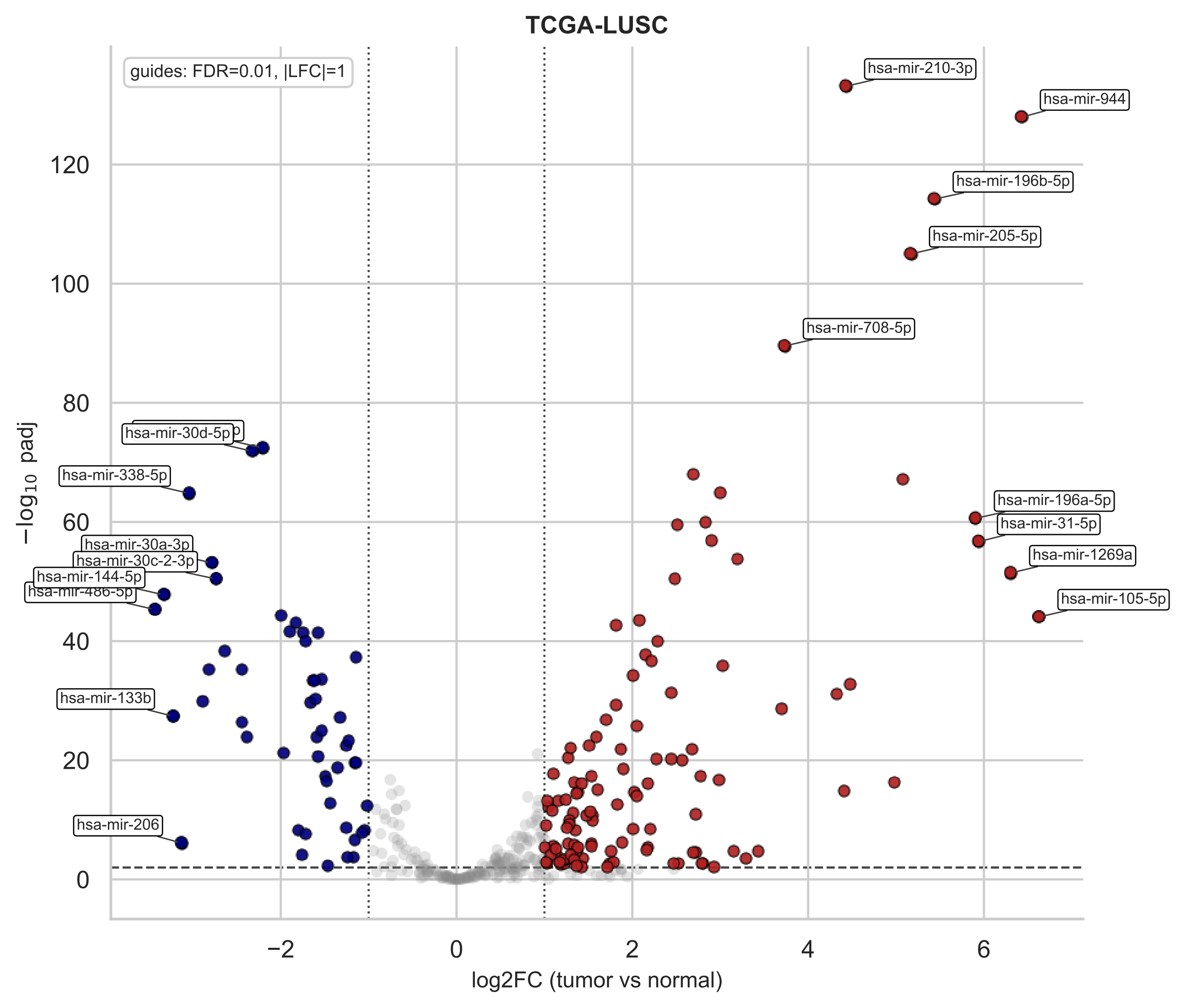
TCGA-LUSC
guides: FDR=0.01, |LFC|=1
hsa-mir-210-3p
hsa-mir-944
hsa-mir-196b-5p
hsa-mir-205-5p
hsa-mir-708-5p
hsa-mir-30d-5p
hsa-mir-338-5p
hsa-mir-196a-5p
hsa-mir-31-5p
hsa-mir-30a-3p
hsa-mir-30c-2-3p
hsa-mir-144-5p
hsa-mir-486-5p
hsa-mir-1269a
hsa-mir-105-5p
hsa-mir-133b
hsa-mir-206
120
100
80
60
40
20
0
−2
0
2
4
6
−log10 padj
log2FC (tumor vs normal)

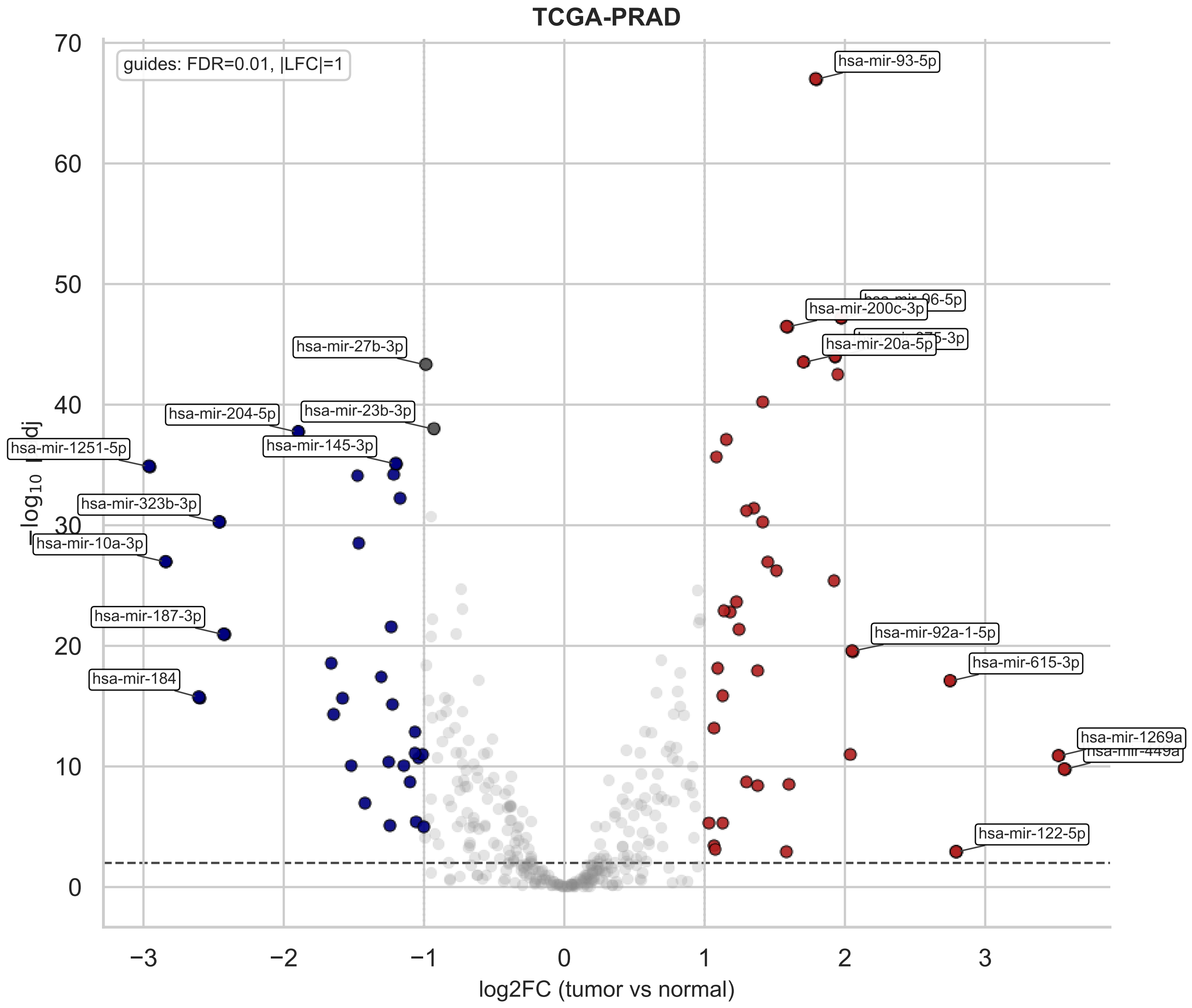

TCGA-PRAD
guides: FDR=0.01, |LFC|=1
hsa-mir-93-5p
hsa-mir-200c-3p
hsa-mir-20a-5p
hsa-mir-27b-3p
hsa-mir-204-5p
hsa-mir-23b-3p
hsa-mir-1251-5p
hsa-mir-145-3p
hsa-mir-323b-3p
hsa-mir-10a-3p
hsa-mir-187-3p
hsa-mir-184
hsa-mir-92a-1-5p
hsa-mir-615-3p
hsa-mir-1269a
hsa-mir-122-5p
log2FC (tumor vs normal)

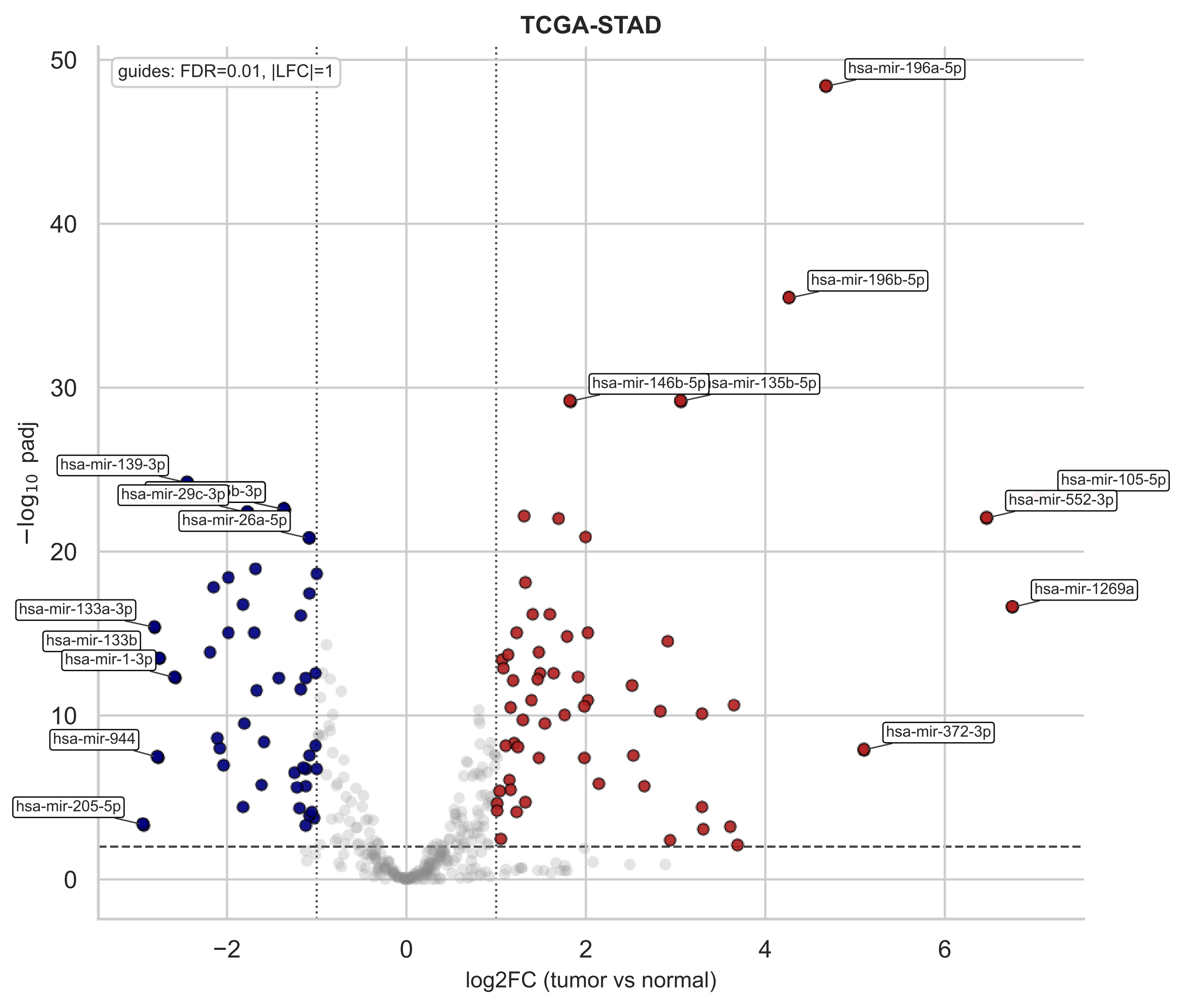

TCGA-STAD
guides: FDR=0.01, |LFC|=1
hsa-mir-196a-5p
hsa-mir-196b-5p
hsa-mir-146b-5p
sa-mir-135b-5p
hsa-mir-105-5p
hsa-mir-552-3p
hsa-mir-1269a
hsa-mir-372-3p
hsa-mir-139-3p
b-3p
hsa-mir-29c-3p
hsa-mir-26a-5p
hsa-mir-133a-3p
hsa-mir-133b
hsa-mir-1-3p
hsa-mir-944
hsa-mir-205-5p
−log10 padj
log2FC (tumor vs normal)
0
10
20
30
40
50
−2
0
2
4
6

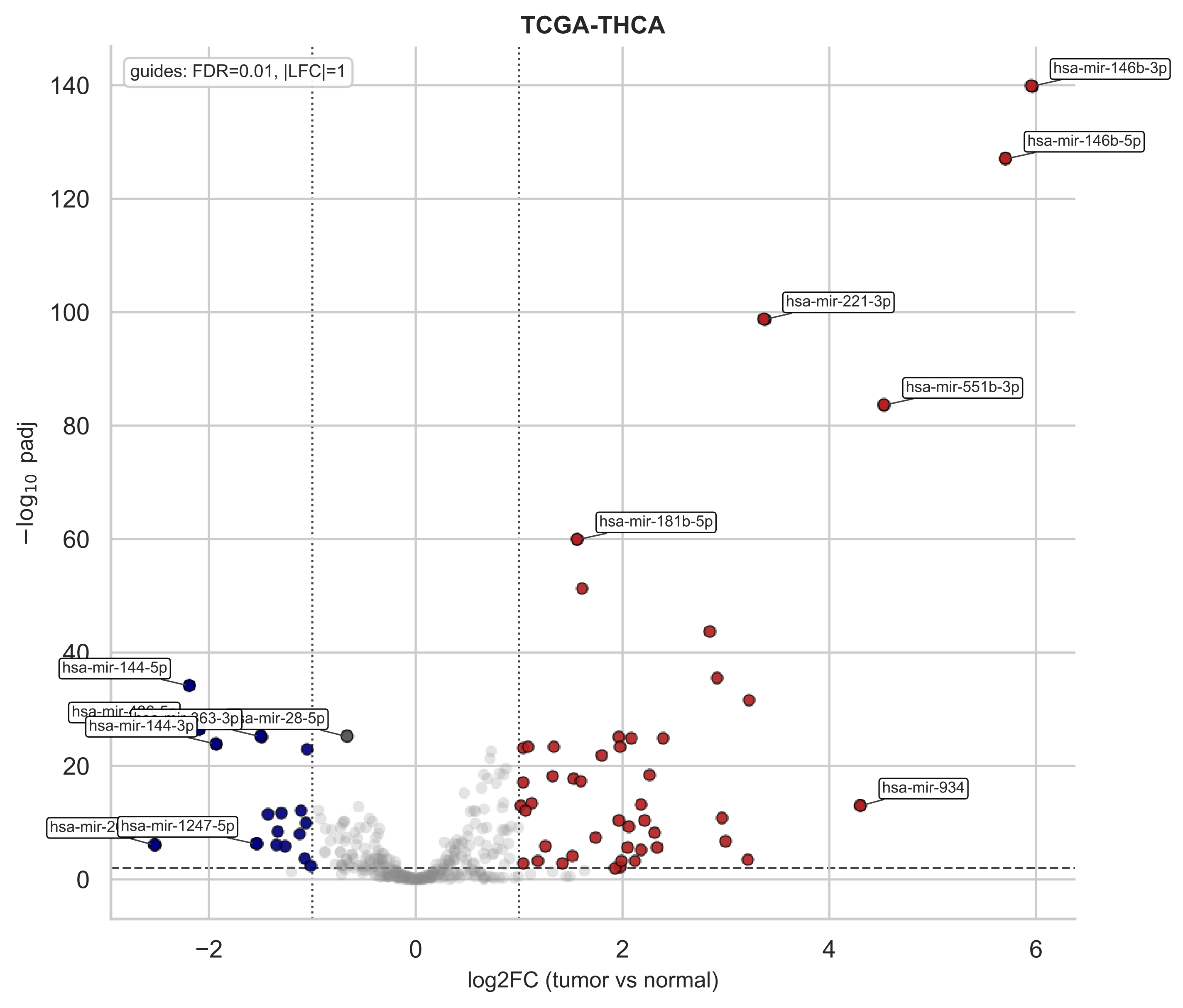

TCGA-THCA
guides: FDR=0.01, |LFC|=1
hsa-mir-146b-3p
hsa-mir-146b-5p
hsa-mir-221-3p
hsa-mir-551b-3p
hsa-mir-181b-5p
hsa-mir-144-5p
hsa-mir-28-5p
hsa-mir-144-3p
hsa-mir-934
hsa-mir-1247-5p
−log10 padj
log2FC (tumor vs normal)

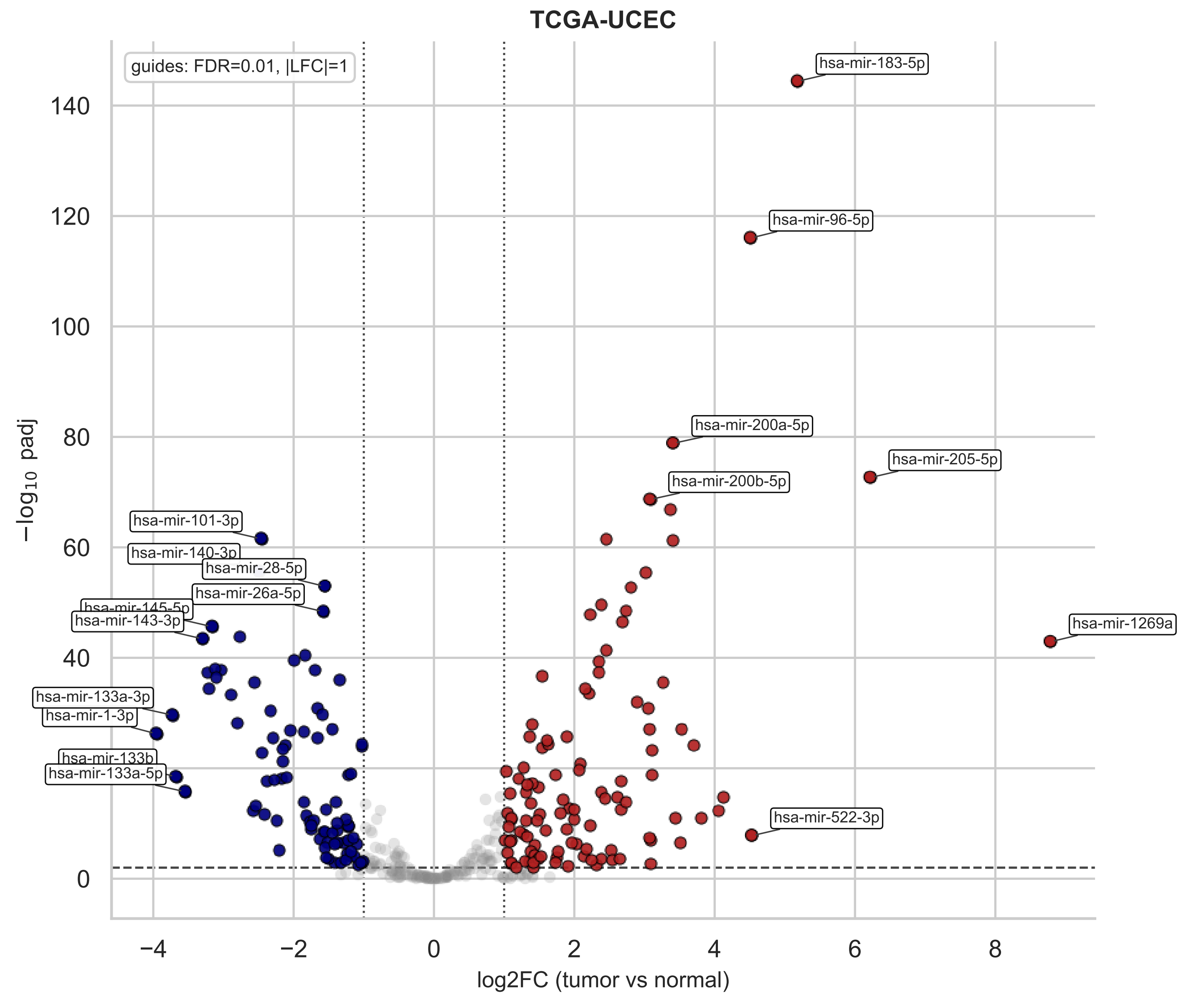
TCGA-UCEC
guides: FDR=0.01, |LFC|=1
hsa-mir-183-5p
hsa-mir-96-5p
hsa-mir-200a-5p
hsa-mir-205-5p
hsa-mir-200b-5p
hsa-mir-101-3p
hsa-mir-140-3p
hsa-mir-28-5p
hsa-mir-26a-5p
hsa-mir-145-5p
hsa-mir-143-3p
hsa-mir-1269a
hsa-mir-133a-3p
hsa-mir-1-3p
hsa-mir-133b
hsa-mir-133a-5p
hsa-mir-522-3p
−log10 padj
log2FC (tumor vs normal)
0
20
40
60
80
100
120
140
−4
−2
0
2
4
6
8
