## Supplementary Figure 2 for "A pan-cancer analysis of microRNA tissue specificity and its association with dysregulation"

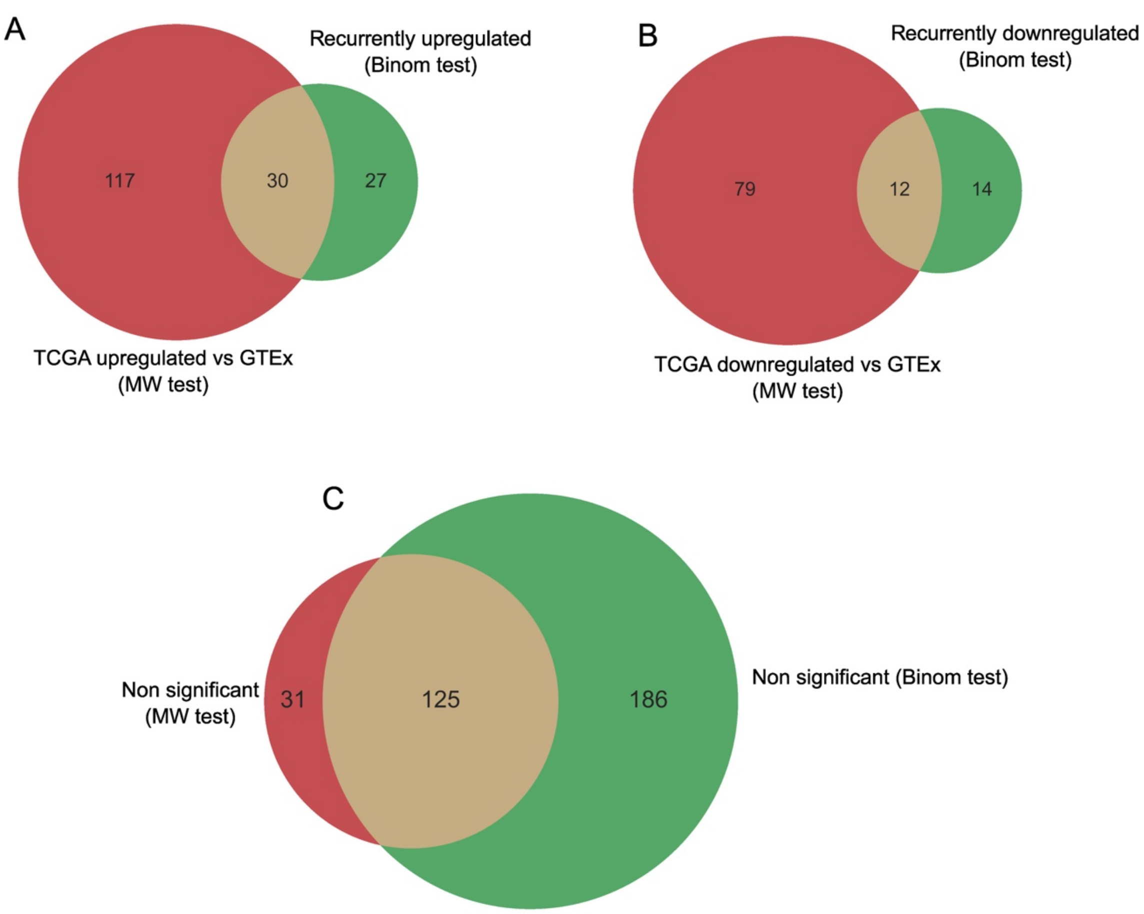


**Supplementary Figure 2.** Overlap between miRNA dysregulation sets identified by two complementary approaches: recurrent dysregulation (binomial test) and the Mann–Whitney (MW) test comparing TCGA and GTEx mean expression levels. Venn diagrams illustrate the intersection of miRNA sets classified by each method as (A) upregulated, (B)downregulated, and (C) non-significantly dysregulated.
